# On islands, evolutionary but not functional originality is rare

**DOI:** 10.1101/822064

**Authors:** S. Veron, R. Pellens, A. Kondratyeva, P. Grandcolas, Rafaël Govaerts, M. Robuchon, T. Haevermans, M. Mouchet

## Abstract

Functionally and evolutionary original species are those whose traits or evolutionary history are shared by few others in a given set. These original species promote ecosystem multifunctionality, the ability to cope with an uncertain future, future benefits to society and therefore have a high conservation value. A potential signal of their extinction risks is their rarity (stating for geographic range-restriction in this study). On islands, life in isolation conducted to the rise of a multitude of original forms and functions as well as to high rates of endemism. Not only patterns and processes of insular originality are unexplained but the relationship between originality and rarity is still unknown. The aim of this study is to assess how original insular species are, to explore whether original species are rare or not and to investigate the factors that may explain the rarity of original species. We first compared the functional and evolutionary originality of monocotyledon species and whether continental or insular species were more original. We found that species restricted to islands were more original than continental species and, although functionally and evolutionary original species were dissimilar, many occurred on similar territories so that regional conservation strategies may allow to conserve these distinct forms. Yet, evolutionary original species were significantly more range-restricted than those which were distinct in their traits. Reflecting their rarity, evolutionary original species had low dispersal abilities and were found on islands where settlement may have been facilitated. On the opposite, functionally original species could reach a wider set of islands by being transported on long-distances. While some mechanisms may both explain rarity and originality such as extinctions, others may be specific to each of these biodiversity facets, in particular diversification, niche shift and expansion, and dispersal power. Implications for conservation are huge: original species are range-restricted and mostly found in the most threatened systems of the world, i.e. islands, endangering the reservoir of features against an uncertain future.

## Introduction

Originality of a given species in a set is the oddity of its biological characteristics, whatever they are, relatively to all other species of the set (Pavoine, Bonsall, Dupaix, Jacob, & Ricotta, 2017). Originality can refer to evolutionary history or to functional traits (e.g. Kondratyeva, Grandcolas & Pavoine, 2019; Pavoine, Ollier, & Dufour, 2005; Pavoine et al., 2017; Redding, Mazel, & Mooers, 2014; Mouillot, Culioli, Pelletier, & Tomasini, 2008; Violle et al., 2017). Originality has been poorly investigated in community ecology or biogeography (but see Mouillot, Graham, Villéger, Mason, & Bellwood, 2013) but its value for biodiversity conservation is well supported. Indeed, in addition to capture significant amount of phylogenetic or functional diversity (Mouillot et al., 2008; Faith, 1992; Kondratyeva et al. 2019), evolutionary and functionally original species may insure key ecosystem processes and provide services to humanity. Functional and evolutionary originality may sometimes be decoupled (Faith, 1992; Pollock, Thuiller, & Jetz, 2017) and should be differentiated due to the distinct benefits they may provide. Regarding measures of functional originality, they are based on a reduced number of traits generally related to ecosystem functioning. For example, (Petchey, Hector, & Gaston, 2004) used 12 plant traits and showed that functionally original species may have a large contribution to ecosystem functions by increasing plant biomass production. Loss of functionally original species may then directly impact ecosystem stability, resilience and multi-functionality (Fonseca & Ganade, 2001; Pendleton, Hoeinghaus, Gomes, & Agostinho, 2014; Bracken & Low, 2012; Oliver et al., 2015). Evolutionary original species may also have a leading contribution to ecosystem functioning because phylogenetic isolation may be an indicator of distinct species ecological roles (Cadotte, Cardinale, & Oakley, 2008; (Redding et al., 2008; Marc W. Cadotte & Jonathan Davies, 2010). Most of all, phylogenetic diversity represents a reservoir of yet-to-be-discovered resources to humanity and evolutionary original species may highly contribute to these option-values (Faith, 2018). Using together functional and evolutionary originality may allow to identify species that capture irreplaceable ecosystem functions and services to humanity. However, functional and evolutionary originality have rarely been studied jointly. Systems in which the study of both biodiversity facets could be highly valuable are islands. Insular systems have a huge importance in biodiversity conservation to preserve the heritage of our planet and they have a leading role of “natural laboratory” for biogeography and evolutionary ecology (Whittaker, Fernández-Palacios, Matthews, Borregaard, & Triantis, 2017; Warren et al., 2015). Islands are home of species which are found nowhere else on Earth and which represent a unique evolutionary history as well as a multitude of forms and functions. Original species may have a high contribution to the island biota, but their role in shaping insular diversity just begin to be investigated (Veron, Haevermans, Govaerts, Mouchet, & Pellens, 2019; Veron et al., 2019). In addition, very little is known about how originality arose and is distributed, so that investigations on this topic may bring insights that go beyond the scope of island biogeography.

A pre-requisite to the joint study of functional and evolutionary originality is to assess how these biodiversity facets are related. As stated above, findings about this issue are contradictory and it has rarely been studied in an insular context. On islands, the relationship between phylogenetic and functional originality may be weak as early diversification and simultaneous cladogenesis resulted in the lack of phylogenetic signal in insular species traits (Losos, 2008). A famous example is the radiation of *Bidens* in Hawaii: closely-related species have greater diversity of traits (e.g. growth form, floral morphology) than in the rest of the world probably due to the diversity of habitat types and to loss of dispersability (Knope, Morden, Funk, & Fukami, 2012). As for Hawaiian *Drosophila*, sexual selection has resulted in clearly distinguishable traits among closely-related (Gillespie & Clague, 2009). In addition, phenotypic character changes of insular species do not necessarily involve speciation or a phylogenetic branching pattern so that variation in evolutionary history do not reflect the variation of some specific traits (Whittaker & Fernández-Palacios, 2007). The first issue we investigated was therefore the extent to which functional and evolutionary originality are related and we specifically tested the assumption that evolutionary and functional originality are less correlated than on the mainland.

Another unresolved question is whether originality is higher on islands in comparison to continental areas. On islands, a long evolutionary history in isolation coupled with unfilled niches and oceanic climate may have given rise to many evolutionary and functionally original species. Extinctions on the continents may also have isolated species in the tree of life and in the functional trait space, creating evolutionary and functionally original insular species (Gillespie & Roderick, 2002; Grandcolas, Nattier, & Trewick, 2014). However, whether insular species are more original than continental ones has been poorly studied so far and maybe highly taxa specific (Jetz et al., 2014). We therefore explored whether insular species are more original than continental ones, which may add a new line of argument for the preservation of insular biodiversity.

Some original species may be at risk due to their rarity (here stating for geographic range-restriction). Range-restriction is indeed a key factor of extinction risks. Many research found that low geographic range size was the main predictor of these risks and it is also one of the main factors of the threat status of species in the IUCN RedList (Rodrigues, Pilgrim, Lamoreux, Hoffmann, & Brooks, 2006; Bland, Collen, Orme, & Bielby, 2015; Veron et al., 2016). Like original species, range-restricted species provide unique functions in an ecosystem as well as unique services to humanity (David Mouillot, Graham, et al., 2013; Leitão et al., 2016). Species which are both rare and original may then support highly vulnerable and unique functions and option-values (Rosauer, Laffan, Crisp, Donnellan, & Cook, 2009; Mouillot, Graham, et al., 2013; Violle et al., 2017; Faith, 2018). Many of such species may be found on islands. Indeed, a remarkable feature of islands is their high rates of endemism which may outcompete these in the mainland by a factor of 9 (Kier et al., 2009). More generally, many insular species are spatially range-restricted and found only on a few islands. Yet, how range-restricted are original species on islands is still poorly known and, as far as we know, has never been tested in the case of functional originality. To fill this gap, we explored the potential risks of losing the most original species by assessing the relationship between species originality and rarity. Beyond practical conservation implications, assessing the relationship between originality and rarity may allow to shed light on the mechanisms shaping both originality and range-restriction. For example, past extinctions may have both isolated species on a phylogenetic tree and/or function space and shrunk species area of distribution. Rarity of original species can also be influenced by their dispersal capacities (Flather et al. 2007). In an insular context, factors influencing colonization, such as the possibility of an island to have been at reach during its geological history and species dispersal traits have a strong influence on the distribution of biodiversity on islands (Gillespie et al., 2012; Weigelt et al., 2015). We therefore investigated how the dispersal capacities of original species and the features of the islands they occur on can give evidence of their rarity.

Originality on islands is still a pristine research field and through our analyses we tackled four main questions that may lay the foundations of the study of originality on islands having implications in both conservation and biogeography 1) Are evolutionary and functionally original species similar on islands? 2) Are insular species more evolutionary and functionally original than mainland species? 3) Are insular original species rare? 4) Can the rarity of original species be attributed to their dispersal capacities and to the biogeographic characteristic of islands? To address these issues we focused on the group of Monocotyledons (Monocots), a morphologically and functionally diverse clade representing a quarter of flowering plant diversity such as all orchids, palms and cereals. The origin of monocot species represents a diversity of evolutionary and ecological processes. Monocots species are also wide-spread across islands and continents, their phylogeny is well-resolved and their traits well-documented. The clade of monocotyledon is therefore a well-suited group for the study of originality on islands.

## Method

### Occurrence data

We used the e-monocot database (emonocot.org) to extract data on both the spatial distribution of monocot species and delimitations of 126 islands (TDWG 3rd level). E-monocot compiles records from several botanical institutions for all 70,000 monocotyledons. We kept only native and terrestrial species from this database.

### Estimating evolutionary and functional originality

We followed results from (Pavoine et al., 2017) to choose the metrics of evolutionary and functional originality (EO and FO, respectively). We followed the recommendations of the authors to use a distance-based metric over dendrogram metrics because the latter may be biased by clustering methods, the use of a low number of traits and of the use of non-ultra-metric trees (Mouchet et al., 2008; Pavoine et al., 2017). We first estimated functional and phylogenetic distances thanks to the Gower’s distance (Legendre & Legendre, 2012) and derived an originality score calculated with the distinctDis function of the package adiv (Pavoine, 2019). EO is estimated in million years of evolution and FO has no units and ranges between 0 and 1.

We estimated monocot FO with six traits compiled by (Díaz et al., 2015), adult plant height, stem specific density, leaf size expressed as leaf area, leaf mass per area, leaf nitrogen content per unit mass, and diaspore mass (see Díaz et al., 2015 for a description). The traits we chose have been widely used and recognized as fundamentally representatives of plant ecological strategies (Díaz et al., 2015). Missing values were imputed by performing random forest algorithm a thousand times, and by then estimating the mean of the 1000 imputation. We performed a sensitivity analysis (Appendix 1) to i. assess the extent to which missing values may have influenced our results ii. estimate the correlation between traits iii. Assess the contribution of each trait to FO scores. We did not include dispersal mode in the measure of functional originality because it was used later as an explanatory factor of the originality-rarity relationship. By doing so we assumed that dispersal mode was more related to the geographic extent of a species than to its function in an ecosystem or its response to environmental conditions.

To estimate EO thanks to the distinctDis function, the Monocot phylogeny we used was built by extracting monocotyledon species from a larger supertree (Qian & Jin, 2016) which is an updated version of a mega-phylogeny of plant species (Zanne et al., 2013). EO was estimated in million years of evolution and FO has no units and range from 0 (lowest originality possible) to 1 (highest originality possible). Due to differences in data availability, and in order to estimate originality on the largest sets of species possible, evolutionary originality was estimated on a set of 6.682 species and FO on a set of 2.281species.

### Are species more original on continents or on islands?

We distributed species among three non-overlapping categories a) insular endemics, i.e. present only on islands; b) continental endemics, i.e. present only on continents; c) insular non-endemic species, i.e. present on both continents and islands. We then estimated the distribution and the average of originality scores among the three categories. We tested whether the average originality was significantly different between the three categories: we randomized species among categories 1000 times, estimated the new average originalities per category in the randomized set and compared them to the observed originalities in each category. A correction under phylogenetic constraint of the average originality score and its significance per category is provided in Appendix 2 but did not influence the results.

### Relationship between Evolutionary and Functional originality

For each category of species, we performed correlation tests between functional and evolutionary originality and measured the phylogenetic signal of each of the six traits individually (Kstar test). To perform these particular analyses, we excluded species that did not have both phylogenetic and functional information. As EO and FO scores are relative to the set of species used to calculate originality, estimating their correlation required to re-calculate a second EO score from the similar set used to estimate FO (i.e. 2.281 species).

Considering again the original set of species (i.e. 6.682 species for EO and 2.281 species for FO), we then only focused on species present in islands (insular endemic and non-endemic species) and drew maps of the number of top 5% original species, i.e. the 5% most original species, in islands both for FO and EO. This included 110 non-endemic species and 40 insular endemics for EO and 38 insular non-endemic and 15 insular endemics for FO.

### Are insular original species geographically rare?

Focusing on species present on islands, i.e. both insular endemics and non-endemics, we tested the correlation (Pearson’s test) between the originality score and *geographic rarity*, measured as *the number of islands each species occur on*. The fewer islands a species occur on the rarer it is. We also used linear regressions corrected for phylogenetic signal in model residuals (Revell, 2010), when necessary. We then specifically tested whether the most original species occurred on fewer islands than expected. We classified species with “top originality”, “high originality”, “moderate originality”, “low originality”, “very low originality” (ranked in the 5%, 25%, 50%, 75% and >75% of the originality score distribution, respectively) and calculated for each class the mean number of islands each species occurred on. We randomized species among classes 1000 times and estimated whether the observed mean number of islands species occurred on per originality class was lower than in the randomized set (see also Appendix 2). Finally, we identified islands where species were both among the top original species and were single endemics (found on a single island). We did not use indices incorporating both originality and geographic rarity as they may sometimes be less relevant for practitioners (Rosauer et al., 2009) and may be difficult to interpret at the functional level.

### Dispersal modes, biogeographic characteristic of islands and species originality

Coupling dispersal features of original insular species and the characteristics of the islands where they occur may help to understand the distribution of original species and possibly their rarity (e.g. (Veron, Haevermans, et al., 2019). First, we compiled information on insular species dispersal mode from (Carvajal-Endara, Hendry, Emery, & Davies, 2017) as well as from the TRY, SID, FRUBASE, BROT and LEDA databases. Dispersal modes corresponded to dispersal by animal, wind, water, unassisted and others (mainly transportation by humans). Species dispersal strategy was then estimated as either long-distance dispersal (animal, wind, water), short distance (unassisted) or unknown (others). We calculated the mean species originality per dispersal mode and strategy and assessed their significance by using null models based on the random distribution of dispersal modes among species. In the case of zoochory, we also explored the effect of the identity of the species dispersing seeds (bird [flying/non flying], mammal [flying/non flying], reptile [terrestrial-marine], insects) on originality (Appendix 3). However, diaspore mass is a trait used in our measure of FO that may be related to dispersal strategy and their association thus needed to be tested. We found that long-distance dispersal was moderately related to heavy diaspore mass (Welch two-sided test p-val=0.07). Moreover, as “diaspore mass” did not prevail on other traits in the estimation of FO scores (Appendix 1), the relationship between FO and dispersal strategy may have been poorly influenced by the moderate association between “diaspore mass” and “dispersal strategy”.

Following our investigation on dispersal strategies of original species, we investigated how the possibility of an island to be/have been at reach may explain the distribution of original species. In particular, we focused on the biogeographic characteristics of islands that may be linked to past, present and future dispersal events (Table 1) although we acknowledge that some of these island features may also drive species diversification and therefore originality.

**Table 1:** Assumptions about how island features can relate to species geographic rarity

| *Island characteristics* | *Assumption about the relation to dispersal potential* | *Reference* |
| --- | --- | --- |
| *Distance to the continent (km)* | An original species having reached a remote island may be a good disperser | Global Island Database (GID; (UNEP-WCM, 2013)) |
| *Proportion of surrounding landmass* | Colonization events may be less frequent on/from isolated islands | (P. Weigelt, Jetz, & Kreft, 2013) |
| *Elevation (m)* | Elevation may provide a climatic refuge for ancient lineages which may consequently rarely disperse | (P. Weigelt et al., 2013) |
| *Area (m<sup>2</sup>)* | Successful colonization events may be more probable towards/from large islands | GID (Depraetere & Dall, 2007) |
| *Oceanic or Continental* | Species present on oceanic islands may occur following colonization and are likely to be relatively good dispersers | (P. Weigelt et al., 2013) |
| *Glaciated or non-Glaciated* | The presence of ancient species on a glaciated island may be due to colonization and tendency to dispersal | Literature; pers. comm. |
| *Age (million years)* | An ancient species occurring on a recent island may reflect colonization, and the probability of the species to be a good disperser may be higher | Literature; pers. com. |
| *Latitude (decimal degrees)* | Variable used to highlight a possible spatial effect | GID (UNEP-WCM, 2013) |
| *Longitude (decimal degrees)* | Variable used to highlight a possible spatial effect | GID (UNEP-WCM, 2013) |

Our aim here was to assess the characteristics of the islands where original species occur, potentially revealing the tendency for species to disperse or not (Table 1). To do so, we estimated for each species the average value of each of the biogeographical features of the islands in which it occurred. Each species was therefore associated to 9 values representing these average island features (Table 1). We then performed generalized linear models and multi-model selection with species originality as the response variable and island average features as the explanatory variables (see also Appendix 2 for models performed under phylogenetic constraint). We identified the strongest interactions thanks to Boosted Regression Trees, which were added in the multi-model selection process.

## Results

### Functional and Evolutionary originality are decoupled

At the exception of diaspore mass of insular species, none plant functional trait exhibited a phylogenetic signal (Table 2). Functional and evolutionary originality were weakly correlated, although the correlation coefficient was higher in species occurring on islands (Pearson’s correlation test: cor=0.16, 0.13, 0.018 and 0.093 in insular endemics, insular non-endemics, continental endemics and all monocot species, respectively). This result was robust to our sensitivity tests (Appendix 1).

**Table 2:**
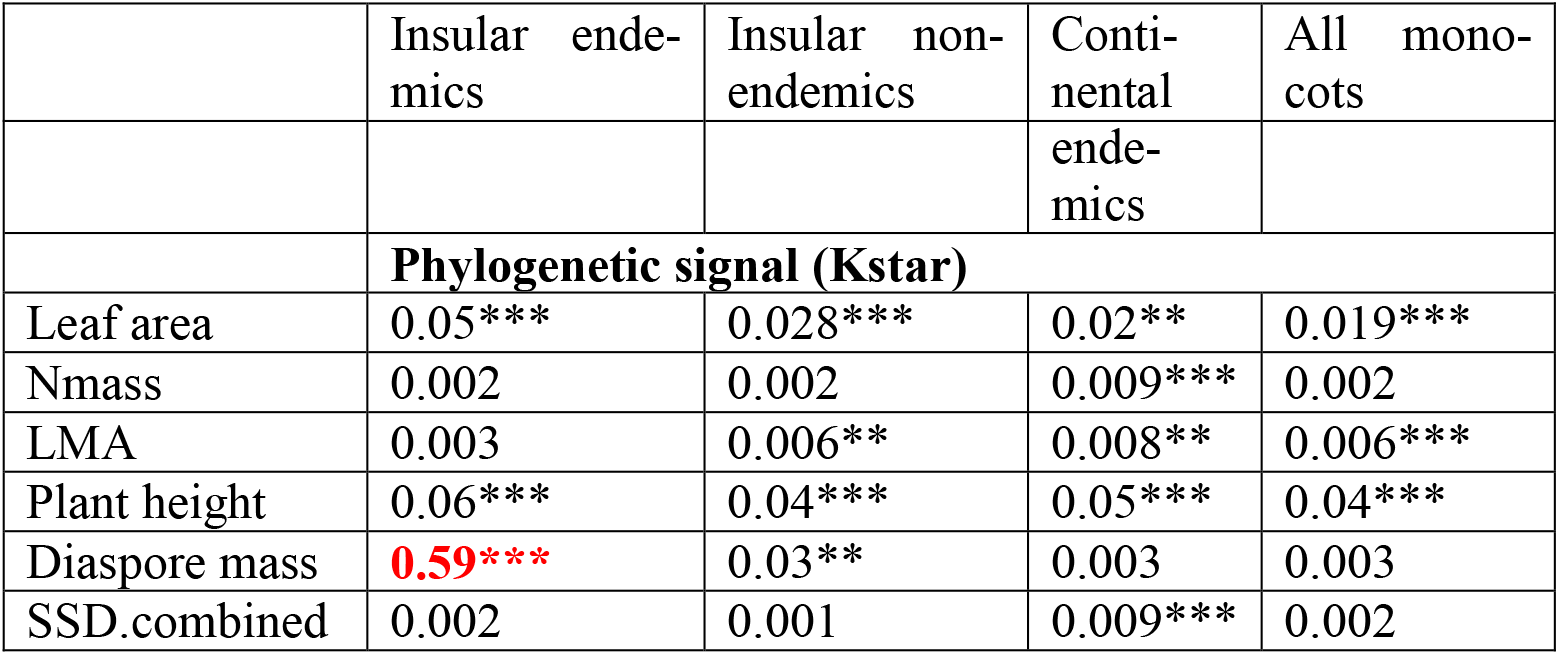
Phylogenetic signal (Kstar) in 6 ecological plant traits. Black values mean there is no phylogenetic signal and bold red values indicate a phylogenetic signal

|  |  |  |  |  |
| --- | --- | --- | --- | --- |
|  | Insular endemics | Insular non-endemics | Continental | All monocots |
|  |  |  | ende-<br>mics |  |
|  | **Phylogenetic signal (Kstar)** |  |  |  |
| Leaf area | 0.05*** | 0.028*** | 0.02** | 0.019*** |
| Nmass | 0.002 | 0.002 | 0.009*** | 0.002 |
| LMA | 0.003 | 0.006** | 0.008** | 0.006*** |
| Plant height | 0.06*** | 0.04*** | 0.05*** | 0.04*** |
| Diaspore mass | **0.59\*\*\*** | 0.03** | 0.003 | 0.003 |
| SSD.combined | 0.002 | 0.001 | 0.009*** | 0.002 |

### Species endemic to islands are more original

Evolutionary originality was estimated on a set of 6.682 species and FO on a set of 2.281species. Islands with the highest number of evolutionary original endemic monocotyledons were Borneo (being home of 10 top 5% original species), the Philippines (9 species), Sumatra (7 species) and Japan (6 species; Figure 1). The maximum concentration of species with top 5% functional original endemic species in a given island was 4 (Philippines), while several other islands were home of 3 top 5% functionally original endemic species (New Guinea, New Zealand, Norfolk Island). Thirteen islands were home of both top 5% functionally original endemic species and top 5% evolutionary original endemic species.

**Figure 1:**
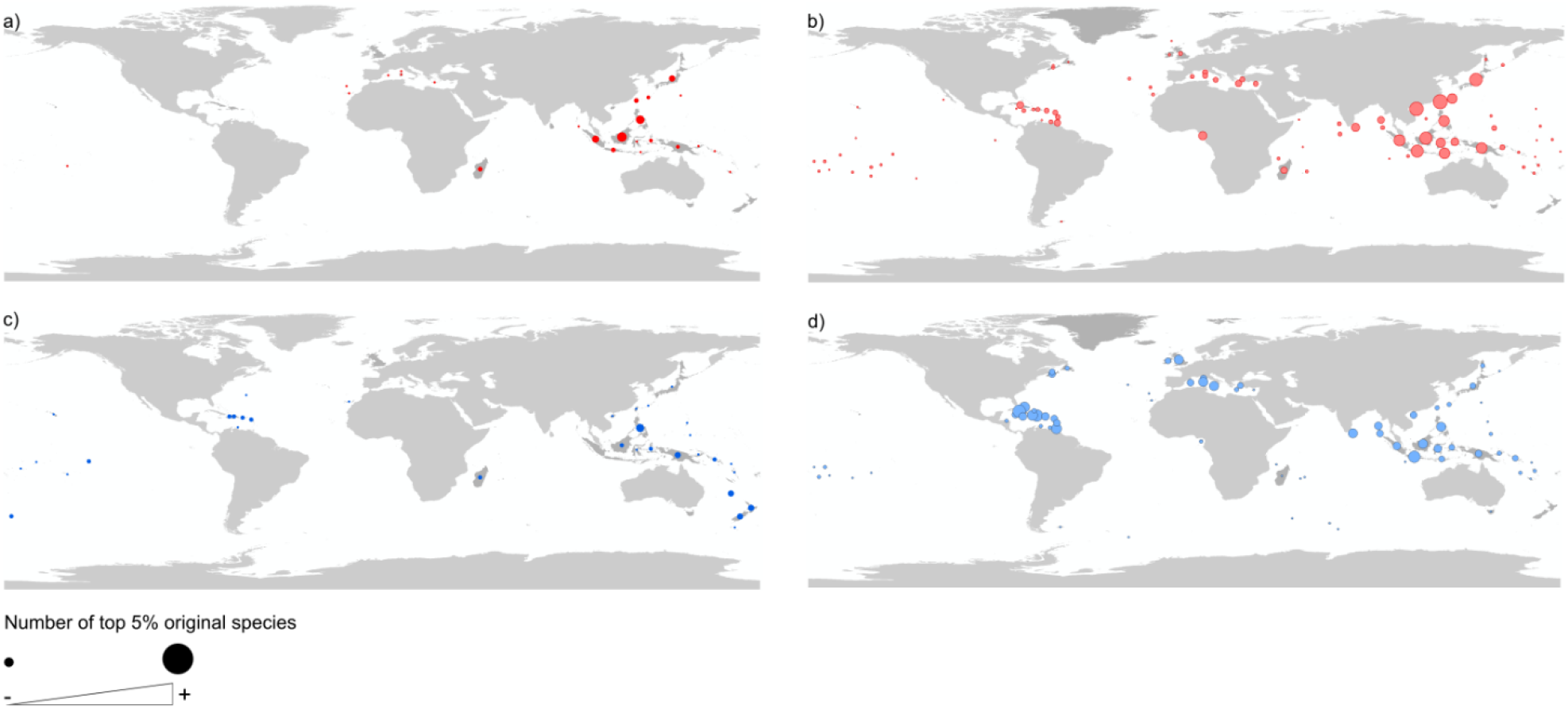
Number of top original species per island for a) EO of endemic insular species b) EO of non-endemic insular species c) FO of endemic insular species d) FO of non-endemic insular species

In non-endemic species, the highest numbers of top 5% evolutionary original species were found in Taiwan (22 species), Hainan (20 species), Japan (17 species), Borneo and Java (16 species). Regarding FO, Cuba was home of 11 top 5% original species, Java and Dominican Republic of 9 species while Hainan, the Bahamas, and Trinidad and Tobago harbored 8 top 5% functionally original species. There were 63 islands where both top 5% evolutionary and functionally non-endemic original species occurred.

By estimating the average originality according to the endemic or non-endemic status of species we found that mean EO and FO was equal to 236.7 and 0.061 for species endemic to islands, 234.9 and 0.043 for non-endemic insular species and 224.5 and 0.047 for continental species only (Figure 2). Average EO was higher than expected for species endemic to islands and those found only in continents. Using phylogenetic analysis of variance models, EO was the highest for insular endemic species (Appendix 2). Average FO was significantly high for insular species whether they were endemic or not.

**Figure 2:**
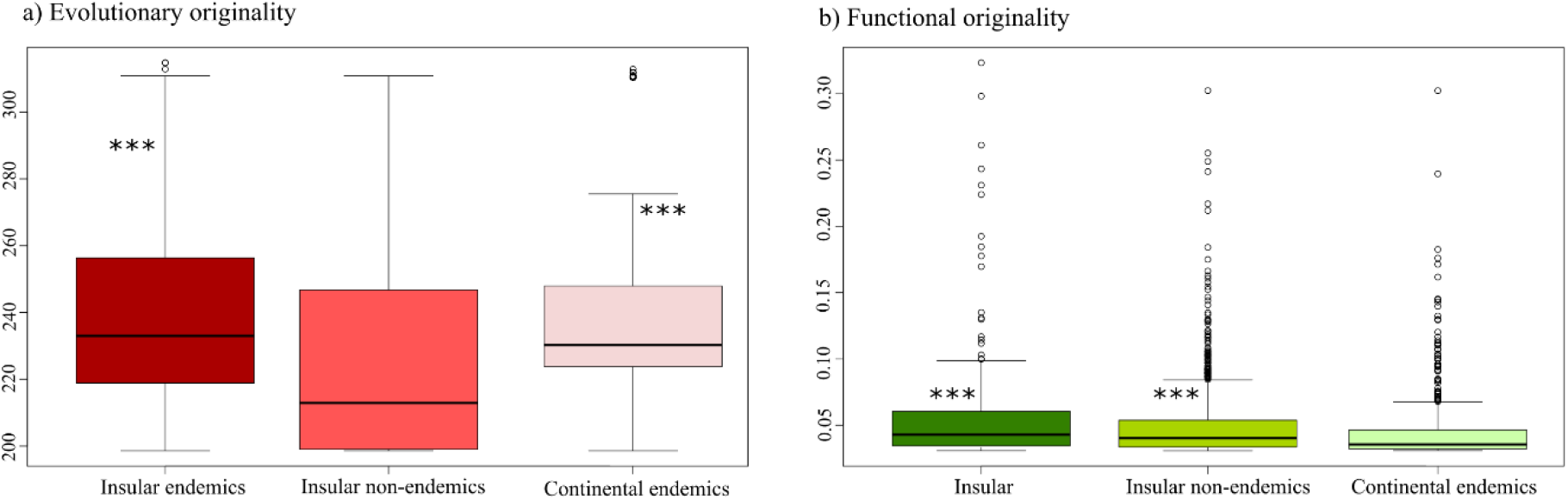
Species originality depending on its origin. *** states for a p-value<0.001

### Evolutionary but not functionally original species are unusually rare

The top 5% original endemic species regarding EO and FO were present on, on average, 1.7 and 3.2 islands, respectively (Table 5). The original non-endemic species occurred on 5.0 to 5.4 islands, on average.

**Table 4:** Effects and importance of island features on EO and FO estimated from multi-model selection

| EO | Species restricted to islands |  |  | Species found on both islands and continents |  |  |
| --- | --- | --- | --- | --- | --- | --- |
|  | *Estimate* | *Pr(>z)* | *Importance* | *Estimate* | *Pr(>z)* | *Importance* |
| **SLMP** | -4.2 | . | 0.62 | -0.34 |  | 0.29 |
| **Distance** | -2.6 | * | 0.89 | -2.21 | *** | 1 |
| **Area** | 0.62 |  | 0.28 | -0.45 |  | 0.32 |
| **Elevation** | -6.3 | *** | 1 | -0.31 |  | 0.29 |
| **Age** | 5.1 | ** | 0.98 | -1.5 | * | 0.88 |
| **GMMC** | 7.07 | *** | 1 | 4.9 | *** | 1 |
| **Glaciated** | -5.9 | *** | 1 | -3.5 | *** | 1 |
| **Latitude** | -6.79 | *** | 1 | -2.6 | ** | 1 |
| **Longitude** | -4.25 | ** | 0.79 | 0.58 |  | 0.35 |
| **Latitude:SLMP** | 7.21 | *** | 1 |  |  |  |
| **Latitude:Age** | 1.53 |  | 0.29 |  |  |  |
| **Distance:Elevation** |  |  |  | 1.5 | . | 0.66 |
| **Glaciated:Distance** |  |  |  | 0.21 |  | 0.24 |

| FO | Species restricted to islands |  |  | Species found on both islands and continents |  |  |
| --- | --- | --- | --- | --- | --- | --- |
|  | *Estimate* | *Pr(>z)* | *Importance* | *Estimate* | *Pr(>z)* | *Importance* |
| **SLMP** | 0.33 | *** | 1 | -0.087 | *** | 1 |
| **Near_dist** | 0.049 | ** | 0.98 | -0.058 | *** | 1 |
| **Area** | -0.083 | *** | 1 | -0.005 |  | 0.39 |
| **Elevation** | 0.029 |  | 0.42 | -0.027 | *** | 1 |
| **Age** | NA |  |  | -0.033 | *** | 1 |
| **GMMC** | -0.22 | *** | 1 | 0.013 | * | 0.85 |
| **Glaciated** | 0.081 | *** | 1 | -0.04 | *** | 1 |
| **Latitude** | -0.26 | *** | 1 | -0.005 |  | 0.27 |
| **Longitude** | 0.084 | *** | 1 | -0.09 | *** | 1 |
| **Latitude:Distance** | 0.01 |  | 0.25 |  |  |  |
| **SLMP:Latitude** | -0.14 | *** | 1 |  |  |  |
| **SLMP:Longitude** |  |  |  | 0.001 |  | 0.25 |
| **SLMP:Elev** |  |  |  | 0.029 | *** | 1 |

**Table 5:** Average geographic range of species, i.e. the number of islands it occurs on, and whether it is lower or larger than expected by chance per category of originality (one-tailed tests). Red means species occur on fewer islands than expected (p-value<0.05). Blue means species occur on more islands than expected (p-value<0.05)

|  | Distribu-<br>tion | Top origi-<br>nal | Very origi-<br>nal | Moderate | Low | Very Low |
| --- | --- | --- | --- | --- | --- | --- |
| Evolution-<br>ary Originality | Insular<br>endemics | Range=1.7 | Range=1.7 | Range=2.1 | Range=2.0 | Range=1.8 |
|  | Insular<br>non-en-<br>demics | Range=5.0 | Range=3.7 | Range=5.9 | Range=7.4 | Range=6.4 |
| Functional<br>Originality | Insular<br>endemics | Range=3.2 | Range=2.4 | Range=1.7 | Range=1.8 | Range=1.8 |
|  | Insular<br>non-en-<br>demics | Range=5.4 | Range=6.4 | Range=7.2 | Range=6.1 | Range=6.0 |

Geographic range had a significant negative effect on originality for EO of insular non-endemic species (Pearson’s test: cor = −0.15 p-value = 4.794e-11) and a significant positive effect on FO of insular endemic species (cor = 0.43 p-value = 3.794e-7). In other cases, no significant effect was found (EO endemics: cor = −0.043 p-value = 0.26; FO non-endemics: cor = −0.028 p-value = 0.28).

Regarding the classification of species according to their originality score, the top 5% EO endemic species on fewer islands than expected by chance while top FO species occurred on more islands than expected (Table 3, Appendix 2). For insular non-endemic species, high EO species also occurred on fewer islands than expected by chance (Table 3, Appendix 2). Conversely, species with low or very low EO occurred on more islands than expected by chance. As for non-endemic FO, those with moderate FO occurred on more islands than expected.

31 species were both endemic to a single island and among the top 5% most evolutionary original species. Among the areas which concentrated the highest number of these rare and original species, Borneo harbor 7 of them, 5 were found in the Philippines, Japan and Sumatra, and 3 in Madagascar (Figure 3). 6 species were single endemics and top 5% functionally original. They occurred on Madagascar (2 species), Lord Howe island (2 species), Bermuda (1 species) and the Canaries (1 species). Regarding non-endemic species 27 were top evolutionary original species and found on a single island. They were mainly concentrated on Equatorial Guinea islands (7 species), Japan (5 species), Cuba (4 species), Hainan (3 species) and Ceylan (2 species). 11 non-endemic species were top functionally original and had a single insular location, in particular Trinidad (3 species), Bali, Tasmania, Sicily, Japan, Equatorial Guinea, Ceylan, Corea, Taiwan (1 species).

**Figure 3.**
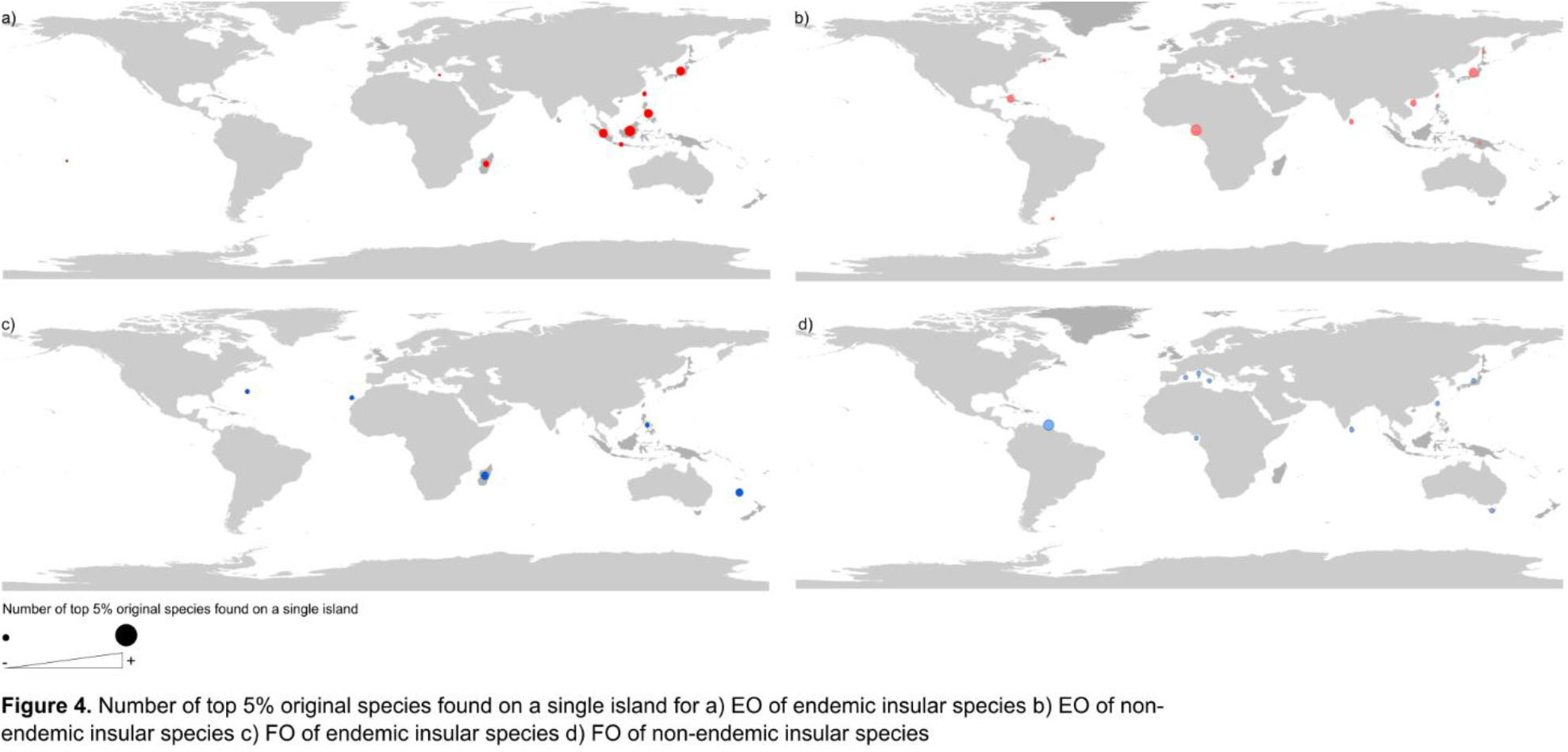
Number of top 5% original species found on a single island for a) EO of endemic insular species b) EO of non-endemic insular species c) FO of endemic insular species d) FO of non-endemic insular species.

### Dispersal abilities of original species

On average, species that dispersed on small distances were more evolutionary original than species dispersing on long-distances (Figure 4a). Endemic evolutionary original species were mainly transported by wind, animals or their mode of dispersal was unassisted. Non-endemic evolutionary original species had mainly an “unassisted” mode of dispersal. Regarding FO, species dispersing on long distances were more functionally original, especially zoochor and anemochor species (Figure 4c and d).

**Figure 4:**
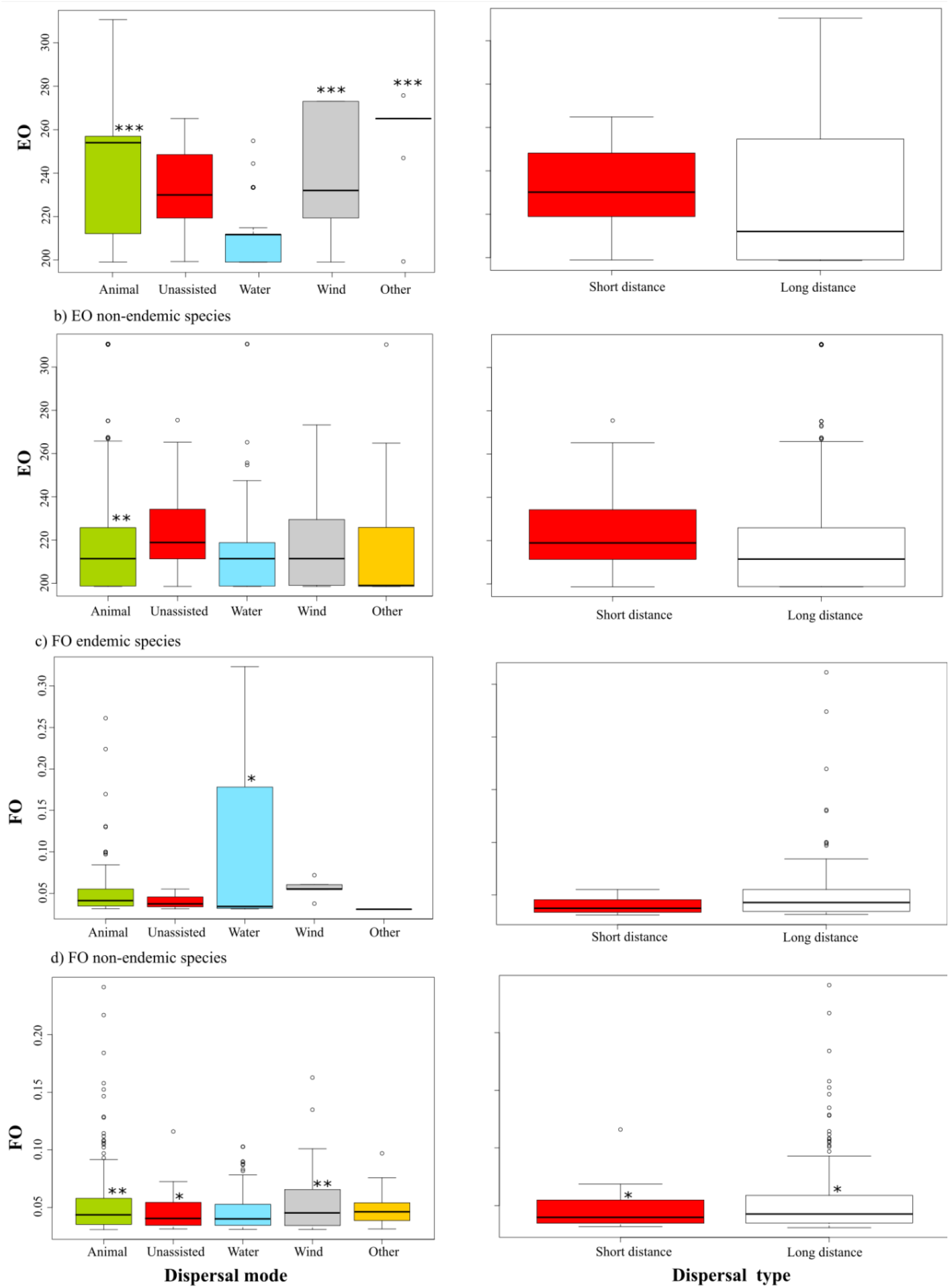
Originality scores depending on the dispersal mode and type. *** means p-value<0.001; ** p-value<0.05; * p-value<0.1

### Possibility of islands to be at reach and distribution of originality

One common point to functionally and evolutionary original species is that they were located at low latitudes, indicating a spatial effect on originality (Table 4). The magnitude of the longitude effect was generally higher for FO than for EO. We observed a high number of differences between the effects of island features on FO and EO. Whatever they were endemic to islands or not, evolutionary original species tended to be found on islands that were continental, non-glaciated, flat and close to the mainland. In addition, endemic evolutionary original species occurred mainly on old islands while non-endemic evolutionary original species were found on more recent islands. The effect of the proportion of surrounding landmass was not significant on the evolutionary originality of non-endemic species and was negative in the case of endemic species but with a low significance (Table 4a). Area had little effect on evolutionary original species whether they were endemic or non-endemic to islands. Regarding FO we found important differences between endemic and non-endemic species. Endemic functionally original species tended to occur on islands that were oceanic, glaciated, small, relatively elevated, close to other landmasses but far from the continent (Table 4b). On the opposite, non-endemic functionally original species were present on islands of continental origin, close to the mainland, and which were relatively young (Table 4b).

## Discussion

### Decoupling of functional and evolutionary originality

In this study we drew a first picture of species originality on islands. Original species have a strong contribution to the uniqueness of island biota, to divergence with the mainland and to centers of endemism (Veron, Haevermans, et al., 2019; Veron, Mouchet, et al., 2019). Among the islands where concentrate both functionally and evolutionary original species are Indonesian islands, the Philippines, Taiwan, Hainan, New-Guinea, Japan, Sumatra, Madagascar. However, some islands harbor only EO species and others only FO species. Importantly, the sets of the most functionally and evolutionary original species were not similar and functional traits of plants representing their ecological strategies had weak phylogenetic signals. There are many possible reasons why FO and EO are not congruent (e.g. Cornwell et al., 2014; Véron, Saito, Padilla-García, Forest, & Bertheau, 2019). On islands, species representative of adaptive radiation, and consequently being closely related and having very close evolutionary originality values, may have very distinct traits in order to fill the niche space let vacant by the low species richness or/and to avoid competition. On the opposite, the process of relictualization, i.e. extinctions of close-relatives on the continent, may have artificially isolated species in the tree-of-life, but some may have actually diversified relatively recently and share several traits with current species (Grandcolas et al., 2014; Whittaker & Fernández-Palacios, 2007). Another reason for this decoupling is convergent evolution which has been reported in multiple lineages on multiple islands (Whittaker et al., 2017). This is the process by which evolutionary unrelated organisms show similar features as a result of natural selection and adaptation under similar environmental constraints (Mazel et al., 2017). Although eco-evolutionary studies would be needed to better understand the evolution of plant species traits in islands, this provided a first evidence that ecological strategies of plants are not conserved in the phylogeny (see also Cornwell et al., 2014).

### Higher originality on islands than on continents

A second important finding was that island endemic species were on average more functionally and evolutionary original than these occurring on continents. A possible explanation is the isolated nature of islands which allows evolution to take its own course giving birth to species evolutionary isolated from all others and/or that harbor unique forms and functions. Higher evolutionary originality on islands compared to the mainland was however unexpected. Indeed, the initial insular pool is generally a set of the continental pool, therefore one could expect that insular species would not be more ancient and evolutionary isolated than continental ones. Moreover, high speciation rates in islands may result in the diversification of many closely-related species which is expected to decrease the average originality of insular pools. Past history in continents may be one explanation of our contradictory finding of higher evolutionary originality on islands than on continents. One main process could be continental extinction: species loss on continents may have pruned the tree of life so that several insular species lost some close-relatives and consequently became evolutionary original and sometimes endemic (Gillespie & Roderick, 2002). High climatic fluctuations and extreme events in the mainland concomitant with a relative stable climate in islands drove extinctions on the continent and persistence on islands, especially in plants (Cronk, 1997; R. J. Whittaker & Fernández-Palacios, 2007 but see Jetz et al., 2014). Climate change in continents may not only have conducted to extinctions but also to diversification resulting from natural selection of the most adapted forms. The result is similar as for extinctions: the evolutionary isolation of insular forms. Besides, not only species loss on the continent but the very high number of past extinctions on islands may have conducted to the evolutionary isolation of island species (Courchamp, Hoffmann, Russell, Leclerc, & Bellard, 2014; Robert J. Whittaker et al., 2017). This process of isolation depends on the phylogenetic signature of extinctions in insular lineages, which remains to be explored, but our result suggest that they have been clustered in species-poor clades which originated in evolutionary original species. A phylogenetic tree incorporating extinct species coupled to the identification of their past distribution may help to disentangle the roles of extinctions and ancient diversification on insular originality.

As for functional originality we also showed that it was higher on islands than on the mainland. Two main mechanisms may have resulted in a higher functional originality on islands: niche expansion and niche shift. Niche expansion relates to low species richness and high disharmony in islands which resulted in a diversity of vacant niches to be colonized by functionally diverse species (R. J. Whittaker & Fernández-Palacios, 2007). For example, (Diamond, 1970) reported that pacific birds could occupy higher altitudinal ranges, a higher diversity of habitats and a higher diversity of vertical foraging ranges compared to their continental counterparts after they colonized species-poor islands in the pacific. Niche shift is the changing biological and ecological characteristics of insular species compared to the mainland (MacArthur & Wilson, 1967; Simberloff, 1974). The persistence of insular species and especially plants may have been conditioned to adaptations to the oceanic climate of islands resulting in the evolution of distinct ecological strategies.

### Range-restriction of original species

The few studies exploring the relationship between evolutionary originality and geographic rarity showed that original species were not more range-restricted than others (Jetz et al., 2014; Thuiller et al., 2015; Stein et al., 2018). Here, although correlation between rarity and EO was generally low, we found that the most evolutionary original species were rare. Some mechanisms generating originality in insular systems also generate endemism, for example adaptation to particular environmental conditions and the function of refuge of many islands. As stated above, island biota has also suffered a number of extinctions and it is likely that human-induced extinctions on islands have both artificially isolated insular species in the Tree of Life and turned multi-endemic species into single endemics (R. J. Whittaker & Fernández-Palacios, 2007). Extinctions may therefore be both responsible for higher originality and higher range-restriction on islands than on continents but, as we will discuss further, it is not the only factor explaining these patterns.

On average, functionally original species were found on only a small number of islands, which seems to be in accordance with studies showing that rare functions are usually supported by rare species (Kunin & Gaston, 1993; Grenié et al., 2018; Violle et al., 2017). Yet, using null models we found that range-restriction of functionally original was not significant, i.e. functionally original monocot species are not rarer than less original ones. This contradicts previous findings that “*species that have low functional redundancy (…) are rarer than expected by chance”* (David Mouillot, Bellwood, et al., 2013; see also Kunin & Gaston, 1993). Violle et al. (2017) recently defined a multifaceted framework of “functional rarity”, linking species’ functional originality and rarity In their framework 12 forms of functional rarity are possible at two spatial scales (local and regional). The less frequent case of few common species with original traits is possible goes with our findings. A possible explanation is that functionally original species have low level of competition with other species so that they can more easily settle and occupy a vacant niche when they disperse towards it.

### Dispersal capacities of EO and FO species

There are many examples of islands where species distribution is due to dispersal power (Whittaker & Fernández-Palacios, 2007; Borda-de-Água et al., 2017). For example, the Krakatau island has been colonized by ferns whose spores can be transported over long-distances whereas other lineages are absent due to their lack of “ocean-going” dispersal mechanisms (Whittaker, Jones, & Partomihardjo, 1997). The modes of dispersal of original species and their association with short and long-distance dispersal strongly support our results on the spatial distribution of these species. Highly evolutionary original species are transported predominantly on short distances and their mode of dispersal is generally “unassisted”, which prevents ocean crossing. More strikingly, the strongest the association between originality and the “unassisted” mode of dispersal, the higher is the correlation between originality and rarity. A possibility explaining the “unassisted” mode of dispersal of original species may be related to the loss of dispersal of plants occurring on islands due to a reduced herbivory pressure and lower competition. Yet, this loss of dispersal syndrome is generally tied to species having recently diversified (Whittaker et al., 2017). Consequently, two assumptions that can be made but need to be tested are, first, that evolutionary original species have low dispersal abilities because they have diversified in recent times but became isolated through relictualization, and, second, that low dispersal ability is a relictual trait rather than a derived one in original species.

As for functionally original species, they were more widespread than non-original ones and this also goes with our findings about their mode of dispersal. Indeed, only few of them had an “unassisted” mode of dispersal. Rather they were transported through zoochory (especially birds) and anemochory and may have been able to colonize several islands isolated from each other. Additional important factors of dispersal include the direction, seasonality, strength of dispersal agents (Gillespie et al., 2012). Yet, our results show that the mode of dispersal and the distance on which seeds can be transported are key determinants of the distribution of original species on islands.

The dispersal strategies of original species can also be seen from the features of the islands where they are found. The most evolutionary original species are often single-endemics and rarely occur on islands where recent and/or long-distance dispersal would be needed to establish: they are uncommon on remote, glaciated, young and oceanic islands. Evolutionary original species also occur on islands surrounded by a relatively low proportion of landmass, decreasing the possibilities of dispersal. On the contrary, functionally original endemic species, which are more widespread, occurred on islands with opposite features and that may only be colonized by species having high dispersal abilities. Together, the dispersal abilities of original species and the characteristics of the islands where they are present are important factors explaining the rarity of plants on islands.

### Conservation

Originality may have an intrinsic value and its conservation is essential to preserve the breadth of functions necessary against future uncertainty and to maintain the reservoir of future benefits to people (Faith, 1992; Violle et al., 2017). If original species go extinct first - and previous considerations shows that this is a very likely assumption - some irreplaceable traits and evolutionary branches would be lost, strongly affecting ecosystem processes and services. Preserving EO species is essential due to their high contribution to “option-values” for humanity and the conservation of FO species is needed to maintain ecosystem functions and processes. This is all the more true on islands as, at least for monocots, the world most original species are insular.

Preserving species evolutionary history has sometimes been assumed to be a proxy for the conservation of species traits. However, several reasons such as convergent evolution, the consideration of a low number of traits, a fast pace of diversification, sexual selection etc. may blur this correlation (Gerhold, Cahill, Winter, Bartish, & Prinzing, 2015; Faith, 2018; Véron, Saito, Padilla-García, Forest, & Bertheau, 2019). In our study we emphasized a lack of phylogenetic signal showing that the originality of plant ecological strategies may not be reflected by their evolutionary relationships. Therefore, a ‘silver-bullet” strategy aiming at the protection of both species functions and evolutionary history may be inadequate. However, in spite of being different, EO and FO species tend to co-occur on many similar islands which give the opportunity for combined conservation strategies that may protect both functional and evolutionary originality. Further research could then look at the overlap between original species range on a given island which may help to prioritize areas for conservation.

A highly important consideration for conservation is that, among endemic species, the most evolutionary original ones are also the rarest. This was also true for non-endemic species but to a lesser degree. Rare and original species may cumulate a diversity of threats that make them at high risks of extinctions (Jono & Pavoine, 2012). First, rarity may be a signal of higher extinction probabilities (e.g. Harnik, Simpson, & Payne, 2012), in particular for insular endemic species for which none population may survive on the mainland. Second, species occurring on islands are among the most vulnerable on Earth due to the combined effects of human-induced threat, species naiveté and low range-size (Leclerc, Courchamp & Bellard,. 2018). Third, rare and original insular species may also be highly vulnerable to external threats such as climate or land-use changes because they have low dispersal abilities and no land at proximity to escape (Veron, Mouchet, Govaerts, Haevermans, & Pellens, 2019). Finally, evolutionary original species may be at higher extinction risks due to increased extinction risks with age (Warren et al., 2018). The most functionally original species were not rarer (geographic rarity) than other species, but they were distributed in a low number of islands. In addition, the rarest functionally and evolutionary original species generally did not occur on the same islands and regional or local conservation plans may not allow to protect rare species which are both distinct in their traits and evolutionary history. Understanding the diversity of current threats to original species and how they will respond to these pressures is key to assess their extinction status and to propose actions directed toward the preservation of these unique species (Forest et al., 2018; Stein et al., 2018).

Finally, as a bridge between the conservation of biodiversity and human cultural heritage, and to conclude with originality, islands are not only places where the most original and threatened species occur but they also concentrate the most original and threatened human languages (Perrault, Farrell, & Davies, 2017)

## REFERENCES

Bland, L. M., Collen, B., Orme, C. D. L., & Bielby, J. (2015). Predicting the conservation status of data-deficient species: Predicting Extinction Risk. Conservation Biology, 29(1), 250–259. https://doi.org/10.1111/cobi.12372

Borda-de-Água, L., Whittaker, R. J., Cardoso, P., Rigal, F., Santos, A. M. C., Amorim, I. R., … Borges, P. A. V. (2017). Dispersal ability determines the scaling properties of species abundance distributions: a case study using arthropods from the Azores. Scientific Reports, 7(1), 3899. https://doi.org/10.1038/s41598-017-04126-5

Bracken, M. E. S., & Low, N. H. N. (2012). Realistic losses of rare species disproportionately impact higher trophic levels: Loss of rare ‘cornerstone’ species. Ecology Letters, 15(5), 461–467. https://doi.org/10.1111/j.1461-0248.2012.01758.x

Cadotte, M. W., Cardinale, B. J., & Oakley, T. H. (2008). Evolutionary history and the effect of biodiversity on plant productivity. Proceedings of the National Academy of Sciences, 105(44), 17012–17017. https://doi.org/10.1073/pnas.0805962105

Cadotte, Marc W., & Jonathan Davies, T. (2010). Rarest of the rare: advances in combining evolutionary distinctiveness and scarcity to inform conservation at biogeographical scales: Conservation phylo-biogeography. Diversity and Distributions, 16(3), 376–385. https://doi.org/10.1111/j.1472-4642.2010.00650.x

Carvajal-Endara, S., Hendry, A. P., Emery, N. C., & Davies, T. J. (2017). Habitat filtering not dispersal limitation shapes oceanic island floras: species assembly of the Galápagos archipelago. Ecology Letters, 20(4), 495–504. https://doi.org/10.1111/ele.12753

Cornwell, W. K., Westoby, M., Falster, D. S., FitzJohn, R. G., O’Meara, B. C., Pennell, M. W., … Zanne, A. E. (2014). Functional distinctiveness of major plant lineages. Journal of Ecology, 102(2), 345–356. https://doi.org/10.1111/1365-2745.12208

Courchamp, F., Hoffmann, B. D., Russell, J. C., Leclerc, C., & Bellard, C. (2014). Climate change, sea-level rise, and conservation: keeping island biodiversity afloat. Trends in Ecology & Evolution, 29(3), 127–130. https://doi.org/10.1016/j.tree.2014.01.001

Cronk, Q. C. B. (1997). Islands: stability, diversity, conservation. Biodiversity & Conservation, 6(3), 477–493. https://doi.org/10.1023/A:1018372910025

Depraetere, C., & Dall, A. (2007). IBPoW Database. A technical note on a global dataset of islands (p. 58).

Diamond, J. M. (1970). Ecological Consequences of Island Colonization by Southwest Pacific Birds, I. Types of Niche Shifts. Proceedings of the National Academy of Sciences, 67(2), 529–536. https://doi.org/10.1073/pnas.67.2.529

Díaz, S., Kattge, J., Cornelissen, J. H. C., Wright, I. J., Lavorel, S., Dray, S., … Gorné, L. D. (2015). The global spectrum of plant form and function. Nature, 529, 167.

Faith, D. P. (1992). Conservation evaluation and phylogenetic diversity. Biological Conservation, 61(1), 110. https://doi.org/10.1016/0006-3207(92)91201-3

Faith, D. P. (2018). Phylogenetic Diversity and Conservation Evaluation: Perspectives on Multiple Values, Indices, and Scales of Application. In R. A. Scherson & D. P. Faith (Eds.), Phylogenetic Diversity: Applications and Challenges in Biodiversity Science (pp. 1–26). https://doi.org/10.1007/978-3-319-93145-6_1

Fonseca, C. R., & Ganade, G. (2001). Species functional redundancy, random extinctions and the stability of ecosystems. Journal of Ecology, 89(1), 118–125. https://doi.org/10.1046/j.1365-2745.2001.00528.x

Forest, F., Moat, J., Baloch, E., Brummitt, N. A., Bachman, S. P., Ickert-Bond, S., … Buerki, S. (2018). Gymnosperms on the EDGE. Scientific Reports, 8(1), 6053. https://doi.org/10.1038/s41598-018-24365-4

Gerhold, P., Cahill, J. F., Winter, M., Bartish, I. V., & Prinzing, A. (2015). Phylogenetic patterns are not proxies of community assembly mechanisms (they are far better). Functional Ecology, 29(5), 600–614. https://doi.org/10.1111/1365-2435.12425

Gillespie, R. G., Baldwin, B. G., Waters, J. M., Fraser, C. I., Nikula, R., & Roderick, G. K. (2012). Longdistance dispersal: a framework for hypothesis testing. Trends in Ecology & Evolution, 27(1), 4756. https://doi.org/10.1016/j.tree.2011.08.009

Gillespie, R. G., & Roderick, G. K. (2002). ARTHROPODS ON ISLANDS: Colonization, Speciation, and Conservation. Annual Review of Entomology, 47(1), 595–632. https://doi.org/10.1146/annurev.ento.47.091201.145244

Grandcolas, P., Nattier, R., & Trewick, S. (2014). Relict species: a relict concept? Trends in Ecology & Evolution, 29(12), 655–663. https://doi.org/10.1016/j.tree.2014.10.002

Grenié, M., Mouillot, D., Villéger, S., Denelle, P., Tucker, C. M., Munoz, F., & Violle, C. (2018). Functional rarity of coral reef fishes at the global scale: Hotspots and challenges for conservation. Biological Conservation, 226, 288–299. https://doi.org/10.1016/j.biocon.2018.08.011

Harnik, P. G., Simpson, C., & Payne, J. L. (2012). Long-term differences in extinction risk among the seven forms of rarity. Proceedings of the Royal Society B: Biological Sciences, 279(1749), 4969–4976. https://doi.org/10.1098/rspb.2012.1902

Jetz, W., Thomas, G. H., Joy, J. B., Redding, D. W., Hartmann, K., & Mooers, A. O. (2014). Global Distribution and Conservation of Evolutionary Distinctness in Birds. Current Biology, 24(9), 919–930. https://doi.org/10.1016/j.cub.2014.03.011

Jono, C. M. A., & Pavoine, S. (2012). Threat Diversity Will Erode Mammalian Phylogenetic Diversity in the Near Future. PLoS ONE, 7(9), e46235. https://doi.org/10.1371/journal.pone.0046235

Kier, G., Kreft, H., Lee, T. M., Jetz, W., Ibisch, P. L., Nowicki, C., … Barthlott, W. (2009). A global assessment of endemism and species richness across island and mainland regions. Proceedings of the National Academy of Sciences, 106(23), 9322–9327. https://doi.org/10.1073/pnas.0810306106

Knope, M. L., Morden, C. W., Funk, V. A., & Fukami, T. (2012). Area and the rapid radiation of Hawaiian Bidens (Asteraceae): Rapid radiation of Hawaiian Bidens. Journal of Biogeography, 39(7), 1206–1216. https://doi.org/10.1111/j.1365-2699.2012.02687.x

Kondratyeva, A., Grandcolas, P., & Pavoine, S. (2019). Reconciling the concepts and measures of diversity, rarity and originality in ecology and evolution. Biological Reviews.

Kunin, W. E., & Gaston, K. J. (1993). The biology of rarity: Patterns, causes and consequences. Trends in Ecology & Evolution, 8(8), 298–301. https://doi.org/10.1016/0169-5347(93)90259-R

Leclerc, C., Courchamp, F., & Bellard, C. (2018). Insular threat associations within taxa worldwide. Scientific reports, 8(1), 6393.

Legendre, P., & Legendre, L. (2012). Numerical ecology (Third English edition). Amsterdam: Elsevier.

Leitão, R. P., Zuanon, J., Villéger, S., Williams, S. E., Baraloto, C., Fortunel, C., … Mouillot, D. (2016). Rare species contribute disproportionately to the functional structure of species assemblages. Proceedings of the Royal Society B: Biological Sciences, 283(1828), 20160084. https://doi.org/10.1098/rspb.2016.0084

Losos, J. B. (2008). Phylogenetic niche conservatism, phylogenetic signal and the relationship between phylogenetic relatedness and ecological similarity among species. Ecology Letters, 11(10), 995–1003. https://doi.org/10.1111/j.1461-0248.2008.01229.x

MacArthur, R. H., & Wilson, E. O. (1967). The theory of island biogeography. Princeton, NJ: Princeton University Press.

Mazel, F., Wüest, R. O., Gueguen, M., Renaud, J., Ficetola, G. F., Lavergne, S., & Thuiller, W. (2017). The Geography of Ecological Niche Evolution in Mammals. Current Biology, 27(9), 1369–1374. https://doi.org/10.1016/j.cub.2017.03.046

Mouchet, M., Guilhaumon, F., Villéger, S., Mason, N. W. H., Tomasini, J.-A., & Mouillot, D. (2008). Towards a consensus for calculating dendrogram-based functional diversity indices. Oikos, 117(5), 794–800. https://doi.org/10.1111/j.0030-1299.2008.16594.x

Mouillot, D., Culioli, J. M., Pelletier, D., & Tomasini, J. A. (2008). Do we protect biological originality in protected areas? A new index and an application to the Bonifacio Strait Natural Reserve. Biological Conservation, 141(6), 1569–1580. https://doi.org/10.1016/j.biocon.2008.04.002

Mouillot, David, Bellwood, D. R., Baraloto, C., Chave, J., Galzin, R., Harmelin-Vivien, M., … Thuiller, W. (2013). Rare Species Support Vulnerable Functions in High-Diversity Ecosystems. PLoS Biology, 11(5), e1001569. https://doi.org/10.1371/journal.pbio.1001569

Mouillot, David, Graham, N. A. J., Villéger, S., Mason, N. W. H., & Bellwood, D. R. (2013). A functional approach reveals community responses to disturbances. Trends in Ecology & Evolution, 28(3), 167–177. https://doi.org/10.1016/j.tree.2012.10.004

Oliver, T. H., Heard, M. S., Isaac, N. J. B., Roy, D. B., Procter, D., Eigenbrod, F., … Bullock, J. M. (2015). Biodiversity and Resilience of Ecosystem Functions. Trends in Ecology & Evolution, 30(11), 673–684. https://doi.org/10.1016/j.tree.2015.08.009

Pavoine, S. (2019). Package “adiv.”

Pavoine, Sandrine, Bonsall, M. B., Dupaix, A., Jacob, U., & Ricotta, C. (2017). From phylogenetic to functional originality: Guide through indices and new developments. Ecological Indicators, 82, 196–205. https://doi.org/10.1016/j.ecolind.2017.06.056

Pavoine, Sandrine, Ollier, S., & Dufour, A.-B. (2005). Is the originality of a species measurable?: Originality of a species. Ecology Letters, 8(6), 579–586. https://doi.org/10.1111/j.1461-0248.2005.00752.x

Pendleton, R. M., Hoeinghaus, D. J., Gomes, L. C., & Agostinho, A. A. (2014). Loss of Rare Fish Species from Tropical Floodplain Food Webs Affects Community Structure and Ecosystem Multifunctionality in a Mesocosm Experiment. PLoS ONE, 9(1), e84568. https://doi.org/10.1371/journal.pone.0084568

Perrault, N., Farrell, M. J., & Davies, T. J. (2017). Tongues on the EDGE: language preservation priorities based on threat and lexical distinctiveness. Royal Society Open Science, 4(12), 171218. https://doi.org/10.1098/rsos.171218

Petchey, O. L., Hector, A., & Gaston, K. J. (2004). HOW DO DIFFERENT MEASURES OF FUNCTIONAL DIVERSITY PERFORM? Ecology, 85(3), 847–857. https://doi.org/10.1890/03-0226

Pollock, L. J., Thuiller, W., & Jetz, W. (2017). Large conservation gains possible for global biodiversity facets. Nature, 546(7656), 141–144. https://doi.org/10.1038/nature22368

Qian, H., & Jin, Y. (2016). An updated megaphylogeny of plants, a tool for generating plant phylogenies and an analysis of phylogenetic community structure. Journal of Plant Ecology, 9(2), 233–239. https://doi.org/10.1093/jpe/rtv047

Redding, D. W., Hartmann, K., Mimoto, A., Bokal, D., DeVos, M., & Mooers, A. Ø. (2008). Evolutionarily distinctive species often capture more phylogenetic diversity than expected. Journal of Theoretical Biology, 251(4), 606–615. https://doi.org/10.1016/j.jtbi.2007.12.006

Redding, D. W., Mazel, F., & Mooers, A. Ø. (2014). Measuring Evolutionary Isolation for Conservation. PLoS ONE, 9(12), e113490. https://doi.org/10.1371/journal.pone.0113490

Revell, L. J. (2010). Phylogenetic signal and linear regression on species data: Phylogenetic regression. Methods in Ecology and Evolution, 1(4), 319–329. https://doi.org/10.1111/j.2041-210X.2010.00044.x

Rodrigues, A., Pilgrim, J., Lamoreux, J., Hoffmann, M., & Brooks, T. (2006). The value of the IUCN Red List for conservation. Trends in Ecology & Evolution, 21(2), 71–76. https://doi.org/10.1016/j.tree.2005.10.010

Rosauer, D., Laffan, S. W., Crisp, M. D., Donnellan, S. C., & Cook, L. G. (2009). Phylogenetic endemism: a new approach for identifying geographical concentrations of evolutionary history. Molecular Ecology, 18(19), 4061–4072. https://doi.org/10.1111/j.1365-294X.2009.04311.x

Simberloff, D. S. (1974). Equilibrium Theory of Island Biogeography and Ecology. Annual Review of Ecology and Systematics, 5(1), 161–182. https://doi.org/10.1146/annurev.es.05.110174.001113

Stein, R. W., Mull, C. G., Kuhn, T. S., Aschliman, N. C., Davidson, L. N. K., Joy, J. B., … Mooers, A. O. (2018). Global priorities for conserving the evolutionary history of sharks, rays and chimaeras. Nature Ecology & Evolution, 2(2), 288–298. https://doi.org/10.1038/s41559-017-0448-4

Thuiller, W., Maiorano, L., Mazel, F., Guilhaumon, F., Ficetola, G. F., Lavergne, S., … Mouillot, D. (2015). Conserving the functional and phylogenetic trees of life of European tetrapods. Philosophical Transactions of the Royal Society B: Biological Sciences, 370(1662), 20140005–20140005. https://doi.org/10.1098/rstb.2014.0005

UNEP-WCM. (2013). Global islands database. United Nation’s Environmental Program, Cambridge, United Kingdom.

Veron, S., Haevermans, T., Govaerts, R., Mouchet, M., & Pellens, R. (2019). Distribution and relative age of endemism across islands worldwide. Scientific Reports, 9(1), 11693. https://doi.org/10.1038/s41598-019-47951-6

Veron, S., Mouchet, M., Govaerts, R., Haevermans, T., & Pellens, R. (2019). Vulnerability to climate change of islands worldwide and its impact on the tree of life. Scientific Reports, 9(1), 14471. https://doi.org/10.1038/s41598-019-51107-x

Veron, S., Mouchet, M., Grandcolas, P., Govaerts, R., Haevermans, T., & Pellens, R. (2019). Tracking the origin of island diversity: insights from divergence with the continental pool in monocots [Preprint]. https://doi.org/10.1101/678300

Veron, S., Penone, C., Clergeau, P., Costa, G. C., Oliveira, B. F., São-Pedro, V. A., & Pavoine, S. (2016). Integrating data-deficient species in analyses of evolutionary history loss. Ecology and Evolution, 6(23), 8502–8514. https://doi.org/10.1002/ece3.2390

Véron, S., Saito, V., Padilla-García, N., Forest, F., & Bertheau, Y. (2019). The Use of Phylogenetic Diversity in Conservation Biology and Community Ecology: A Common Base but Different Approaches. The Quarterly Review of Biology, 94(2), 123–148. https://doi.org/10.1086/703580

Violle, C., Thuiller, W., Mouquet, N., Munoz, F., Kraft, N. J. B., Cadotte, M. W., … Mouillot, D. (2017). Functional Rarity: The Ecology of Outliers. Trends in Ecology & Evolution, 32(5), 356–367. https://doi.org/10.1016/j.tree.2017.02.002

Warren, B. H., Hagen, O., Gerber, F., Thébaud, C., Paradis, E., & Conti, E. (2018). Evaluating alternative explanations for an association of extinction risk and evolutionary uniqueness in multiple insular lineages: EXTINCTION AND EVOLUTIONARY UNIQUENESS. Evolution, 72(10), 2005–2024. https://doi.org/10.1111/evo.13582

Warren, B. H., Simberloff, D., Ricklefs, R. E., Aguilée, R., Condamine, F. L., Gravel, D., … Thébaud, C. (2015). Islands as model systems in ecology and evolution: prospects fifty years after MacArthur-Wilson. Ecology Letters, 18(2), 200–217. https://doi.org/10.1111/ele.12398

Weigelt, P., Jetz, W., & Kreft, H. (2013). Bioclimatic and physical characterization of the world’s islands. Proceedings of the National Academy of Sciences, 110(38), 15307–15312. https://doi.org/10.1073/pnas.1306309110

Weigelt, Patrick, Daniel Kissling, W., Kisel, Y., Fritz, S. A., Karger, D. N., Kessler, M., … Kreft, H. (2015). Global patterns and drivers of phylogenetic structure in island floras. Scientific Reports, 5(1). https://doi.org/10.1038/srep12213

Whittaker, R. J., & Fernández-Palacios, J. M. (2007). Island biogeography: ecology, evolution, and conservation (Oxford University Press).

Whittaker, R. J., Jones, S. H., & Partomihardjo, T. (1997). The rebuilding of an isolated rain forest assemblage: how disharmonicis the flora of Krakatau?. Biodiversity and Conservation, 6, 1671–1696.

Whittaker, Robert J., Fernández-Palacios, J. M., Matthews, T. J., Borregaard, M. K., & Triantis, K. A. (2017). Island biogeography: Taking the long view of nature’s laboratories. Science, 357(6354), eaam8326. https://doi.org/10.1126/science.aam8326

Zanne, A. E., Tank, D. C., Cornwell, W. K., Eastman, J. M., Smith, S. A., FitzJohn, R. G., … Beaulieu, J. M. (2013). Three keys to the radiation of angiosperms into freezing environments. Nature, 506, 89.

