## Appendix 1 for "On islands, evolutionary but not functional originality is rare"

**Appendix 1: Sensitivity analysis**

a) **Missing trait values**
i) *Correlation between functional originality scores calculated when missing values were imputed or not*

| Trait | Proportion of missing values |
| --- | --- |
| Adult plant height | 46% |
| Leaf size | 68% |
| Stem specific density | 81% |
| Leaf mass per area | 68% |
| Leaf nitrogen content | 75% |
| Diaspore mass | 23% |

**Table 1.** Proportion of missing values per trait

Missing values were imputed by performing random forest algorithm (missForest package, Stekhoven and Buehlmann, 2012) a thousand times, and the mean of the 1000 imputations was estimated. To show how this imputation method may affect our originality results we tested the correlation between functional originality scores when missing trait values were considered as « Non-Attributed » and functional originality scores computed when missing values were imputed. We found that the correlation was positive but moderate (Spearman correlation coefficient r = 0.65, p-value < 0.001). This result was expected, because random forest method imputes missing values for a species according to the non-missing values of other traits.

ii) *Correlation between the number of missing values and functional originality score*

Second, we tested the correlation between the number of species’ missing trait data and its functional originality score. We found that the number of species’ missing trait values had a negative but poor influence on its functional originality score (Spearman’s correlation coefficient r = -0.14, p-value < 0.001). This only relatively supports the expectancy that random forest imputations would smooth trait values and therefore conduct to an under-estimation of the functional originality scores of species with many missing trait values.

iii) *Correlation between functional originality scores calculated with a complete trait data and when missing trait values were created artificially*

Third, we artificially created missing values for species with complete data for all traits (259 species). We then imputed these artificially missing values with a random forest method. We then calculated functional originality scores on the complete set of species and estimated their correlation with the functional originality scores coming from artificially created trait data. First, all values of a given trait were considered as missing. Second, we randomly created 2/3 of missing values among all traits and repeated the randomization a hundred times. Results showed high correlations between functional originality scores computed on a complete dataset and obtained after artificial deletion and imputation of trait values (Table 2).

|  | Leaf area | Leaf Nitrogen content | Leaf Mass per area | Plant height | Diaspore mass | Stem specific density | 2/3 of missing values randomly distributed |
| --- | --- | --- | --- | --- | --- | --- | --- |
| Pearson coefficient | 0.999 | 0.907 | 0.993 | 0.941 | 0.999 | 0.879 | 0.41 |

**Table 2.** Correlation between observed functional originality scores of species with no missing values and functional originality scores obtained after artificially creating and imputing missing values in a particular trait or in all traits at random.

iv) *Relationship between functional and evolutionary originality scores in the set of species having no missing values*

We calculated functional and evolutionary originality scores on the subset of 259 species for which all trait values were available. We found that the correlation estimated on this particular species subset was low (Spearman’s correlation coefficients r = 0.29, p-values < 0.001).

We finally estimated the phylogenetic signal of each trait from complete trait data subset using a Kstar method. At the exception of the trait “Diaspore mass” none trait displayed a phylogenetic signal.

|  | Leaf area | Leaf Nitrogen content | Leaf Mass per area | Plant height | Diaspore mass | Stem specific density |
| --- | --- | --- | --- | --- | --- | --- |
| Kstar | 0.093 p=0.001 | 0.0018  p=0.743 | 0.003  p=0.605 | 0.112  p=0.006 | 2.05  p=0.001 | 0.0036  p=0.396 |

**Table 3.** Phylogenetic signal of each trait under study calculated with Kstar method.

b) **Correlations between traits**

The highest correlation was between N mass and Stem Specific Density (cor=0.65), for other traits correlation was low.

|  | Leaf Area | N mass | LMA | Plant.Height | Diaspore.Mass (mg) | SSD combined |
| --- | --- | --- | --- | --- | --- | --- |
| Leaf.Area | 1 | 0.034 | 0.49 | 0.44 | 0.09 | 0.22 |
| Nmass | 0.034 | 1 | 0.47 | 0.039 | 0.001 | 0.65 |
| LMA | 0.49 | 0.47 | 1 | 0.24 | 0.081 | 0.43 |
| Plant.Height | 0.44 | 0.039 | 0.24 | 1 | 0.33 | 0.51 |
| Diaspore.Mass (mg) | 0.09 | 0.0013 | 0.08 | 0.33 | 1 | 0.16 |
| SSD.combined | 0.22 | 0.65 | 0.43 | 0.51 | 0.16 | 1 |

**Table 4**. Pearson’s correlation coefficient between trait values

c) **Contribution of each trait to the functional originality score**

To test the influence of each trait on species originality we performed a sensitivity analysis by excluding each trait separately and recalculating functional originality with the five remaining traits. Pearson’s correlation coefficient between functional originality calculated on 6 traits and calculated on 5 traits ranged from 0.91 to 0.99. We concluded that none of traits prevailed over others in the calculation of species functional originality.

| Trait excluded | Pearson’s correlation with all-trait functional originality |
| --- | --- |
| Adult plant height | 0.98*** |
| Leaf size | 0.95*** |
| stem specific density | 0.98*** |
| Leaf mass per area | 0.91*** |
| Leaf nitrogen content | 0.99*** |
| Diaspore mass | 0.96*** |

**Table 5**: Correlations between the functional originality score calculated with all traits and

obtained when a given trait was excluded. *** significance level for p-value < 0.001.

d) **Sensitivity to the metric used to calculate functional originality**

To test the effect of choice of the originality metric we calculated functional originality with Fair Proportion index (FP, Isaac, 2007) on species differences represented by a functional dendrogram (hierarchical clustering method h-clust) and found a significant but moderate correlation with functional originality calculated with AVerage index (AV, Pavoine et al., 2017) on species trait-based dissimilarity matrix. We also calculated evolutionary originality with the same indices (FP and AV) respectively on phylogenetic tree and phylogenetic distances in dissimilarity matrix (Pearson’s correlation for evolutionary originality r = 0.45, p-value < 0.001; for functional originality r = 0.40, p-value < 0.001).
