## Appendix 2 for "On islands, evolutionary but not functional originality is rare"

**Appendix : Phylogenetically corrected models**

1. ***Species endemic to islands are more original***

*Method: Phylogenetic Analysis of variance* (package phytools, Revell 2018)

1.1 *Evolutionary Originality*

**ANOVA table: Phylogenetic ANOVA**

Pairwise posthoc test using method = "holm"

Pairwise t-values:

|  | **Insular endemics** | **Continental endemics** | **Insular non-endemics** |
| --- | --- | --- | --- |
| **Insular endemics** |  | 2.14 | 12.72 |
| **Continental enemics** | -2.14 |  | 16.26 |
| **Insular non-endemics** | -12.72 | -16.26 |  |

Pairwise corrected P-values:

|  | **Insular endemics** | **Continental endemics** | **Insular non-endemics** |
| --- | --- | --- | --- |
| **Insular endemics** |  | 0.38 | 0.003 |
| **Continental enemics** | 0.38 |  | 0.003 |
| **Insular non-endemics** | 0.003 | 0.003 |  |

1.2 *Functional Originality*

Pairwise t-values:

|  | **Insular endemics** | **Continental endemics** | **Insular non-endemics** |
| --- | --- | --- | --- |
| **Insular endemics** |  | 7.95 | 6.24 |
| **Continental enemics** | -7.95 |  | -3.9 |
| **Insular non-endemics** | -6.24 | 3.9 |  |

Pairwise corrected P-values:

|  | **Insular endemics** | **Continental endemics** | **Insular non-endemics** |
| --- | --- | --- | --- |
| **Insular endemics** |  | 0.012 | 0.07 |
| **Continental enemics** | 0.012 |  | 0.647 |
| **Insular non-endemics** | 0.07 | 0.647 |  |

2. ***Original species are more range-restricted***
   2.1 Relationship rarity and originality

*Method: Generalized least squares under phylogenetic constraint*

2.1.1 Insular endemics

2.1.1.1 Evolutionary Originality

| **AIC** | **BIC** | **logLik** |
| --- | --- | --- |
| 2094 | 2107 | -1044.15 |

|  | **Value** | **Std.Err** | **t-value** | **P_val** |
| --- | --- | --- | --- | --- |
| **Intercept** | 277.13 | 1.49 | 169.23 | 0.00 |
| **Number of islands** | -0.00038 | 0.008 | -0.04 | 0.97 |

2.1.1.2 Functional Originality

|  | **Value** | **Std.Err** | **t-value** | **P_val** |
| --- | --- | --- | --- | --- |
| **Intercept** | 277.13 | 1.49 | 169.23 | 0.00 |
| **Number of islands** | -0.00038 | 0.008 | -0.04 | 0.97 |

2.1.2 Insular non-endemics

2.1.2.1 Evolutionary originality

|  | **Value** | **Std.Err** | **t-value** | **P_val** |
| --- | --- | --- | --- | --- |
| **Intercept** | 0.000 | 0.0002 | -0.0025 | 0.99 |
| **Number of islands** | -0.00024 | 0.007 | -0.03 | 0.97 |

2.1.2.2 Functional originality

|  | **Value** | **Std.Err** | **t-value** | **P_val** |
| --- | --- | --- | --- | --- |
| **Intercept** | 0.023 | 0.084 | 0.27 | 0.78 |
| **Number of islands** | 0.014 | 0.0012 | 11.48 | 0.0 |

2.2 Rarity in function of the category of evolutionary originality

*Method: Phylogenetic Analysis of variance*

2.2.1 Insular endemics

2.2.1.1 Evolutionary Originality

The tables show the correlations between the range restriction of species belonging to different originality “categories”

Pairwise posthoc test using method = "holm"

**Pairwise t-values:**

|  | **High originality** | **Low originality** | **Moderate originality** | **Very high originality** | **Very low originality** |
| --- | --- | --- | --- | --- | --- |
| **High originality** | 0.0 | -1.32 | -1.69 | 0.72 | -0.08 |
| **Low originality** | 1.32 | 0.0 | -0.39 | 1.65 | 1.39 |
| **Moderate originality** | 1.68 | 0.39 | 0.0 | 1.89 | 1.80 |
| **Very high originality** | -0.72 | -1.65 | -1.89 | 0.0 | -0.81 |
| **Very low originality** | 0.08 | -1.39 | -1.80 | 0.81 | 0.0 |

Pairwise corrected P-values:

|  | **High originality** | **Low originality** | **Moderate originality** | **Very high originality** | **Very low originality** |
| --- | --- | --- | --- | --- | --- |
| **High originality** | 1 | 1 | 1 | 1 | 1 |
| **Low originality** | 1 | 1 | 1 | 1 | 1 |
| **Moderate originality** | 1 | 1 | 1 | 1 | 1 |
| **Very high originality** | 1 | 1 | 1 | 1 | 1 |
| **Very low originality** | 1 | 1 | 1 | 1 | 1 |

2.2.1.2 Functional Originality

|  | **High originality** | **Low originality** | **Moderate originality** | **Very high originality** | **Very low originality** |
| --- | --- | --- | --- | --- | --- |
| **High originality** |  | 1.18 | 1.50 | -2.56 | 1.06 |
| **Low originality** | -1.18 |  | 0.32 | -3.48 | -0.06 |
| **Moderate originality** | -1.50 | -0.32 |  | -3.7 | 0.37 |
| **Very high originality** | 2.56 | 3.48 | 3.74 |  | 3.32 |
| **Very low originality** | -1.06 | 0.061 | 0.37 | -3.31 |  |

Pairwise corrected P-values:

|  | **High originality** | **Low originality** | **Moderate originality** | **Very high originality** | **Very low originality** |
| --- | --- | --- | --- | --- | --- |
| **High originality** | 1 | 1 | 1 | 1 | 1 |
| **Low originality** | 1 | 1 | 1 | 1 | 1 |
| **Moderate originality** | 1 | 1 | 1 | 1 | 1 |
| **Very high originality** | 1 | 1 | 1 | 1 | 1 |
| **Very low originality** | 1 | 1 | 1 | 1 | 1 |

2.2.2 Insular non-endemics

2.2.2.1 Evolutionary Originality

Pairwise posthoc test using method = "holm"

Pairwise t-values:

|  | **High originality** | **Low originality** | **Moderate originality** | **Very high originality** | **Very low originality** |
| --- | --- | --- | --- | --- | --- |
| **High originality** | 0.0 | -7.6 | -4.24 | -2.14 | -6.22 |
| **Low originality** | 7.60 | 0.0 | 3.72 | 2.24 | 1.42 |
| **Moderate originality** | 4.24 | -3.72 | 0.0 | 0.28 | -2.24 |
| **Very high originality** | 2.14 | -2.28 | -0.28 | 0.0 | -1.49 |
| **Very low originality** | 6.22 | -1.44 | 2.24 | 1.49 | 0.0 |

Pairwise corrected P-values:

|  | **High originality** | **Low originality** | **Moderate originality** | **Very high originality** | **Very low originality** |
| --- | --- | --- | --- | --- | --- |
| **High originality** | 1 | 1 | 1 | 1 | 1 |
| **Low originality** | 1 | 1 | 1 | 1 | 1 |
| **Moderate originality** | 1 | 1 | 1 | 1 | 1 |
| **Very high originality** | 1 | 1 | 1 | 1 | 1 |
| **Very low originality** | 1 | 1 | 1 | 1 | 1 |

2.2.2.1 Functional Originality

Pairwise t-values:

|  | **High originality** | **Low originality** | **Moderate originality** | **Very high originality** | **Very low originality** |
| --- | --- | --- | --- | --- | --- |
| **High originality** |  | 0.15 | -1.7 | 1.25 | -0.41 |
| **Low originality** | -0.15 |  | -1.95 | 1.19 | 0.57 |
| **Moderate originality** | 1.69 | 1.95 |  | 2.18 | 1.22 |
| **Very high originality** | -1.25 | -1.19 | -2.18 |  | -1.47 |
| **Very low originality** | 0.41 | 0.58 | -1.22 | 1.48 |  |

Pairwise corrected P-values:

|  | **High originality** | **Low originality** | **Moderate originality** | **Very high originality** | **Very low originality** |
| --- | --- | --- | --- | --- | --- |
| **High originality** | 1 | 1 | 1 | 1 | 1 |
| **Low originality** | 1 | 1 | 1 | 1 | 1 |
| **Moderate originality** | 1 | 1 | 1 | 1 | 1 |
| **Very high originality** | 1 | 1 | 1 | 1 | 1 |
| **Very low originality** | 1 | 1 | 1 | 1 | 1 |

3. ***Dispersal mode***

*Method: Phylogenetic Analysis of variance*

3.1 ***Type of dispersal***
    3.1.1 Insular endemics
          3.1.1.1 Evolutionary originality

Pairwise t-values

|  | ***Short distance*** | ***Long distance*** |
| --- | --- | --- |
| ***Short distance*** |  | -0.47 |
| ***Long distance*** | 0.47 |  |

Pairwise corrected P-values:

|  | ***Short distance*** | ***Long distance*** |
| --- | --- | --- |
| ***Short distance*** |  | 0.76 |
| ***Long distance*** | 0.76 |  |

3.1.1.2 Functional originality

|  | ***Short distance*** | ***Long distance*** |
| --- | --- | --- |
| ***Short distance*** |  | -.1.15 |
| ***Long distance*** | 1.15 |  |

Pairwise corrected P-values:

|  | ***Short distance*** | ***Long distance*** |
| --- | --- | --- |
| ***Short distance*** |  | 0.31 |
| ***Long distance*** | 0.31 |  |

3.1.2 Insular non-endemics

3.1.2.1 Evolutionary Originality

Pairwise t-values

|  | ***Short distance*** | ***Long distance*** |
| --- | --- | --- |
| ***Short distance*** |  | 0.64 |
| ***Long distance*** | -0.64 |  |

Pairwise corrected P-values:

|  | ***Short distance*** | ***Long distance*** |
| --- | --- | --- |
| ***Short distance*** |  | 0.84 |
| ***Long distance*** | 0.84 |  |

3.1.2.2 Functional Originality

Pairwise t-values

|  | ***Short distance*** | ***Long distance*** |
| --- | --- | --- |
| ***Short distance*** |  | -1.44 |
| ***Long distance*** | 1.44 |  |

Pairwise corrected P-values:

|  | ***Short distance*** | ***Long distance*** |
| --- | --- | --- |
| ***Short distance*** |  | 0.41 |
| ***Long distance*** | 0.41 |  |

3.2 ***Mode of dispersal***

*Method: Phylogenetic Analysis of variance*

3.2.1 Insular endemics

3.2.1.1 Evolutionary originality

Pairwise t-values

|  | ***Animal*** | ***Other*** | ***Unassisted*** | ***Water*** | ***Wind*** |
| --- | --- | --- | --- | --- | --- |
| ***Animal*** | 0 | 2.36 | 0.92 | 1.64 | 1.45 |
| ***Other*** | -2.36 | 0 | -1.52 | -0.98 | -1.31 |
| ***Unassisted*** | -0.92 | 1.52 | 0 | 0.63 | 0.33 |
| ***Water*** | -1.64 | 0.98 | -0.63 | 0 | -0.34 |
| ***Wind*** | -1.45 | 1.31 | -0.33 | 0.34 | 0 |

Pairwise corrected P-values

|  | ***Animal*** | ***Other*** | ***Unassisted*** | ***Water*** | ***Wind*** |
| --- | --- | --- | --- | --- | --- |
| ***Animal*** | 0 | 1 | 1 | 0.48 | 1 |
| ***Other*** | 1 | 0 | 1 | 1 | 1 |
| ***Unassisted*** | 1 | 1 | 0 | 1 | 1 |
| ***Water*** | 0.48 | 1 | 1 | 0 | 1 |
| ***Wind*** | 1 | 1 | 1 | 1 | 0 |

3.2.1.1 Functional originality

Pairwise t-values

|  | ***Animal*** | ***Other*** | ***Unassisted*** | ***Water*** | ***Wind*** |
| --- | --- | --- | --- | --- | --- |
| ***Animal*** | 0 | 1.37 | 1.14 | -0.69 | 0.56 |
| ***Other*** | -1.38 | 0 | -0.19 | -1.63 | -0.62 |
| ***Unassisted*** | -1.13 | 0.19 | 0 | -1.43 | -0.43 |
| ***Water*** | 0.69 | 1.64 | 1.44 | 0 | 0.98 |
| ***Wind*** | -0.57 | 0.62 | 0.44 | -0.97 | 0 |

Pairwise corrected P-values

|  | ***Animal*** | ***Other*** | ***Unassisted*** | ***Water*** | ***Wind*** |
| --- | --- | --- | --- | --- | --- |
| ***Animal*** | 0 | 1 | 1 | 1 | 1 |
| ***Other*** | 1 | 0 | 1 | 1 | 1 |
| ***Unassisted*** | 1 | 1 | 0 | 1 | 1 |
| ***Water*** | 1 | 1 | 1 | 0 | 1 |
| ***Wind*** | 1 | 1 | 1 | 1 | 0 |

3.2.2 Insular non-endemics

3.2.2.1 Evolutionary Originality

Pairwise t-values

|  | ***Animal*** | ***Other*** | ***Unassisted*** | ***Water*** | ***Wind*** |
| --- | --- | --- | --- | --- | --- |
| ***Animal*** | 0 | 1.40 | -0.32 | 1.08 | 0.50 |
| ***Other*** | -1.40 | 0 | -1.35 | -0.5 | -0.82 |
| ***Unassisted*** | 0.32 | 1.35 | 0 | 1.048 | 0.65 |
| ***Water*** | -1.08 | 0.54 | -1.04 | 0 | -0.38 |
| ***Wind*** | -0.50 | 0.82 | -0.65 | 0.38 | 0 |

Pairwise corrected P-values

|  | ***Animal*** | ***Other*** | ***Unassisted*** | ***Water*** | ***Wind*** |
| --- | --- | --- | --- | --- | --- |
| ***Animal*** | 0 | 1 | 1 | 0.48 | 1 |
| ***Other*** | 1 | 0 | 1 | 1 | 1 |
| ***Unassisted*** | 1 | 1 | 0 | 1 | 1 |
| ***Water*** | 0.48 | 1 | 1 | 0 | 1 |
| ***Wind*** | 1 | 1 | 1 | 1 | 0 |

3.2.2.2 Functional Originality

Pairwise t-values

|  | ***Animal*** | ***Other*** | ***Unassisted*** | ***Water*** | ***Wind*** |
| --- | --- | --- | --- | --- | --- |
| ***Animal*** | 0 | -0.016 | 1.84 | 1.93 | 0.28 |
| ***Other*** | 0.016 | 0 | 1.48 | 1.38 | 0.22 |
| ***Unassisted*** | -1.84 | -1.48 | 0 | -0.35 | -1.36 |
| ***Water*** | -1.93 | -1.38 | 0.35 | 0 | -1.25 |
| ***Wind*** | -0.28 | -0.22 | 1.36 | 1.25 | 0 |

Pairwise corrected P-values

|  | ***Animal*** | ***Other*** | ***Unassisted*** | ***Water*** | ***Wind*** |
| --- | --- | --- | --- | --- | --- |
| ***Animal*** | 0 | 1 | 1 | 1 | 1 |
| ***Other*** | 1 | 0 | 1 | 1 | 1 |
| ***Unassisted*** | 1 | 1 | 0 | 1 | 1 |
| ***Water*** | 1 | 1 | 1 | 0 | 1 |
| ***Wind*** | 1 | 1 | 1 | 1 | 0 |

4 ***Features of islands where original species are found***

*Method: generalized least squares under phylogenetic constraint*

4.1 Insular endemics

4.1.1 Evolutionary Originality

*Coefficients*

|  | ***Value*** | ***Std.Error*** | ***t-value*** | ***p-value*** |
| --- | --- | --- | --- | --- |
| ***Intercept*** | 275.81 | 1.83 | 150.52 | 0,00 |
| ***Dist*** | -0.002 | 0.017 | -0.12 | 0.89 |
| ***Area*** | 0.006 | 0.042 | 0.15 | 0.87 |
| ***Latitude*** | -0.007 | 0.068 | -0.10 | 0.91 |
| ***Longitude*** | 0.0067 | 0.053 | 0.12 | 0.89 |
| ***SLMP*** | 0.0055 | 0.080 | 0.0686 | 0.94 |
| ***GMMC*** | 0.0060 | 0.037 | 0.16 | 0.87 |
| ***Elev*** | -0.0089 | 0.045 | -0.19 | 0.84 |
| ***Agemax*** | 0.0026 | 0.065 | 0.041 | 0.96 |
| ***Glaciated*** | 0.00074 | 0.015 | 0.047 | 0.96 |

4.1.2 Functional originality

|  | ***Value*** | ***Std.Error*** | ***t-value*** | ***p-value*** |
| --- | --- | --- | --- | --- |
| ***Intercept*** | 0.05372446 | 0.094 | 0.56 | 0.57 |
| ***Distance*** | -0.014 | 0.0075 | -1.98 | 0.049 |
| ***Area*** | 0.042 | 0.010 | 4.05 | 0.0001 |
| ***Latitude*** | 0.016 | 0.010 | 1.63 | 0.10 |
| ***Longitude*** | -0.0032 | 0.011 | -0.28 | 0.77 |
| ***SLMP*** | -0.042 | 0.020 | -2.02 | 0.045 |
| ***GMMC*** | -0.0018 | 0.012 | -0.14 | 0.88 |
| ***Elevation*** | -0.042 | 0.014 | -2.89 | 0.0047 |
| ***Age*** | -0.042 | 0.017 | -2.4 | 0.015 |
| ***Glaciated*** | -0.011 | 0.0031 | -3.75 | 0.0003 |

4.2 Insular non-endemics

4.2.1 Evolutionary originality

*Coefficients*

|  | ***Value*** | ***Std.Error*** | ***t-value*** | ***p-value*** |
| --- | --- | --- | --- | --- |
| ***Intercept*** | 269.35 | 1.02 | 263.37 | 0 |
| ***Distance*** | 0.00004 | 0.0039 | 0.010 | 0.99 |
| ***Area*** | 0.00004 | 0.0037 | 0.0098 | 0.99 |
| ***Latitude*** | -0.0002 | 0.0074 | -0.026 | 0.97 |
| ***Longitude*** | 0.00012 | 0.0062 | 0.019 | 0.98 |
| ***SLMP*** | 0.00009 | 0.0075 | 0.011 | 0.99 |
| ***GMMC*** | 0.00007 | 0.0052 | 0.014 | 0.98 |
| ***Elevation*** | 0.00005 | 0.0043 | 0.011 | 0.99 |
| ***Age*** | -0.00005 | 0.0042 | -0.012 | 0.99 |
| ***Glaciated*** | -0.00011 | 0.0072 | -0.015 | 0.98 |

***4.2.2*** Functional originality

|  | ***Value*** | ***Std.Error*** | ***t-value*** | ***p-value*** |
| --- | --- | --- | --- | --- |
| ***Intercept*** | 0.062 | 0.10 | 0.60 | 0.54 |
| ***Distance*** | 0.0035 | 0.00045 | 7.80 | 0.00 |
| ***Area*** | -0.0019 | 0.00049 | -3.98 | 0.0001 |
| ***Latitude*** | 0.0041 | 0.00090 | 4.56 | 0.00 |
| ***Longitude*** | 0.00129 | 0.00073 | 1.69 | 0.090 |
| ***SLMP*** | 0.0094 | 0.00087 | 10.83 | 0.00 |
| ***GMMC*** | 0.00082 | 0.00062 | 1.31 | 0.18 |
| ***Elevation*** | 0.0027 | 0.00048 | 5.6 | 0.00 |
| ***Age*** | -0.00013 | 0.00049 | -0.27 | 0.78 |
| ***Glaciated*** | -0.0042 | 0.00084 | -5.09 | 0.00 |
