## Appendix 3 for "On islands, evolutionary but not functional originality is rare"

Originality scores depending on dispersal vectors of zoochor species

### a) Endemic species

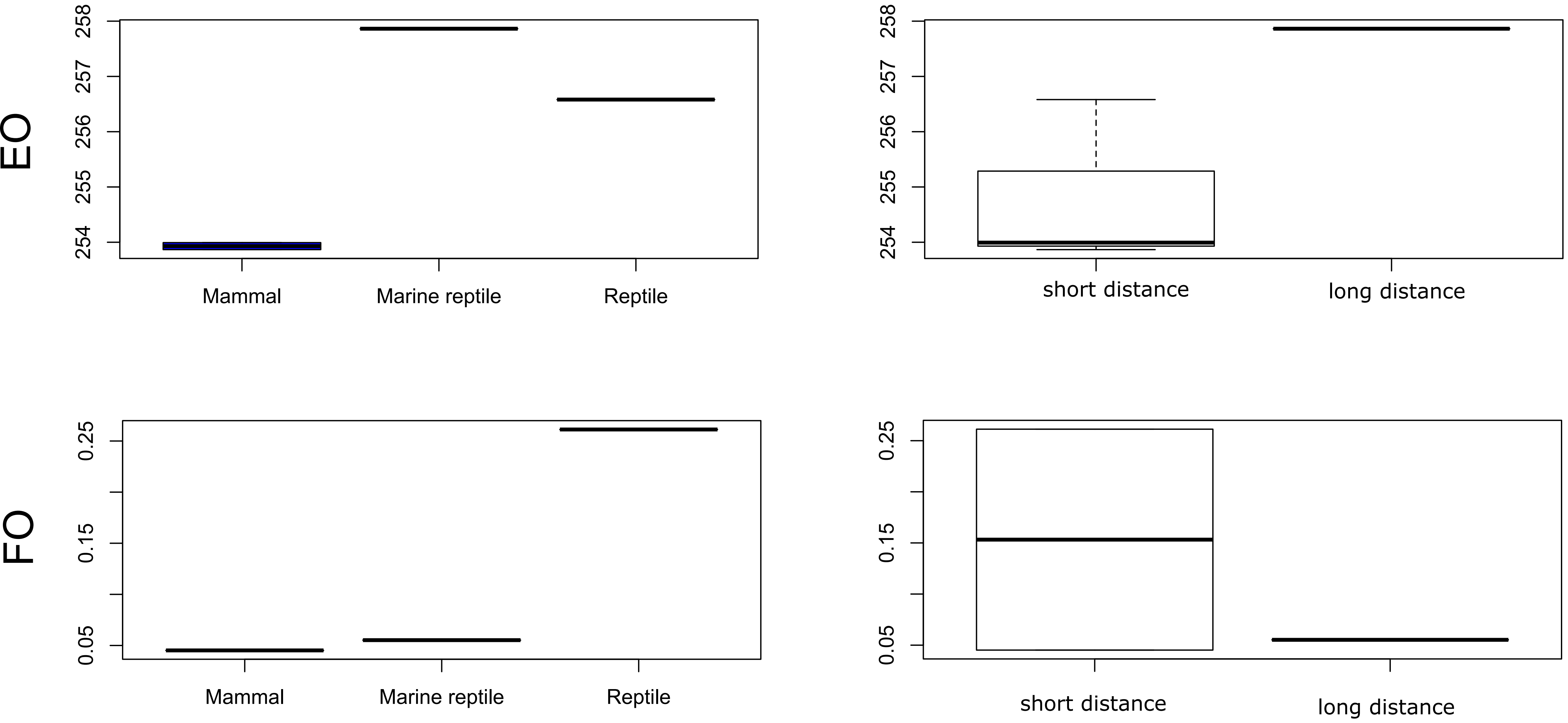


### b) Non endemic species

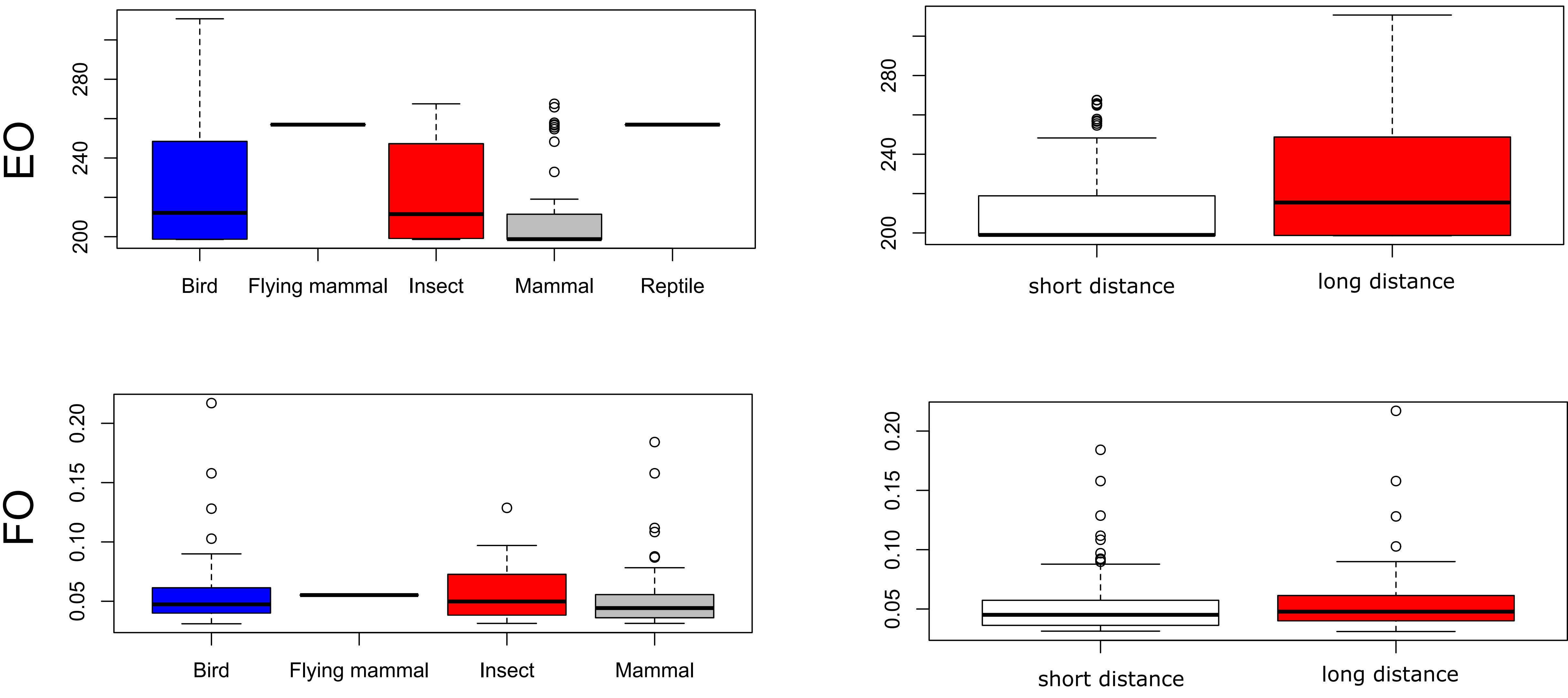
